# HOPS: a quantitative score reveals pervasive horizontal pleiotropy in human genetic variation is driven by extreme polygenicity of human traits and diseases

**DOI:** 10.1101/311332

**Authors:** Daniel M. Jordan, Marie Verbanck, Ron Do

## Abstract

Horizontal pleiotropy, where one variant has independent effects on multiple traits, is important for our understanding of the genetic architecture of human phenotypes. We develop a method to quantify horizontal pleiotropy using genome-wide association summary statistics and apply it to 372 heritable phenotypes measured in 361,194 UK Biobank individuals. Horizontal pleiotropy is pervasive throughout the human genome, prominent among highly polygenic phenotypes, and enriched in active regulatory regions. Our results highlight the central role horizontal pleiotropy plays in the genetic architecture of human phenotypes. The HOrizontal Pleiotropy Score (HOPS) method is available on Github at https://github.com/rondolab/HOPS.

## Background

The term “pleiotropy” refers to a single genetic variant having multiple distinct phenotypic effects. In general terms, the existence and extent of pleiotropy has far-reaching implications on our understanding of how genotypes map to phenotypes (1), of the genetic architectures of traits (2,3), of the biology underlying common diseases (4) and of the dynamics of natural selection (5). However, beyond this general idea of the importance of pleiotropy, it quickly becomes difficult to discuss in specifics, because of the difficulty in defining what counts as a direct causal effect and what counts as a separate phenotypic effect.

One particularly important dividing line in these conflicting definitions is the distinction between vertical pleiotropy and horizontal pleiotropy (6,7). When a genetic variant has a phenotypic effect that then has its own downstream effects in turn, that variant exhibits “vertical” pleiotropy.

For example, a variant that increases low density lipoprotein (LDL) cholesterol might also have an additional corresponding effect on coronary artery disease risk due to the causal relationship between these two traits, thus exhibiting vertical pleiotropy. Vertical pleiotropy has been conceptualized and measured by explicit genetic methods like Mendelian randomization.

In contrast, a genetic cause that directly influences multiple traits, without one trait being mediated by another, would exhibit “horizontal” pleiotropy. Horizontal pleiotropy contains some conceptual difficulties, and consequently can be difficult to measure. In principle, we might imagine selecting a variant and counting how many phenotypes are associated with it. Indeed, several versions of this analysis have been performed for different lists of traits (8,2,3,9). However, the results of these analyses are highly dependent on the exact list of traits used, and traits of interest to researchers previously tend to involve only a small number of phenotypes and/or be heavily biased towards a small set of disease-relevant biological systems and processes. Due to these limitations, it is unknown to what extent horizontal pleiotropy affects genetic variation in the human genome at the genome-wide level.

The proliferation of data sources like large-scale biobanks and metabolomics data that include a wide array of phenotypes in one dataset, combined with the growing public availability of genome-wide association studies (GWASs) summary statistic data, especially for extremely large meta-analyses, has allowed the development of methods that use these summary statistics to gain insight into human biology, and particularly into the genetic architecture of complex traits and diseases (10).

Here, we present the HOrizontal Pleiotropy Score (HOPS) method to measure horizontal pleiotropy using publicly available GWAS summary statistics. We focus on measuring horizontal pleiotropy of SNVs on observable traits, meaning a scenario where a single SNV affects multiple phenotypes without relying on a detectable causal relationship between those phenotypes. Using this framework, we are able to score each SNV in the human genome for horizontal pleiotropy, giving us broad insight into the genetic architecture of pleiotropy. Because our framework explicitly removes correlations between the input phenotypes, and because these phenotypes represent a diverse array of traits and diseases, these insights are largely robust to the specific list of traits studied, and pertain to human biology overall rather than relationships between specific traits.

## Results

### Defining pleiotropy

We narrowly define the scope of pleiotropy as applying only to genetic variants, and particularly variants investigated as part of GWASs. As effects, we are considering phenotypic outcomes measured by GWASs. By our definition, then, pleiotropy means that one variant shows significant associations across GWASs of multiple traits. We additionally restrict the scope of pleiotropy we are considering to include only horizontal pleiotropy, and to exclude vertical pleiotropy (**Figure 1**). To elaborate on this distinction, suppose we have identified a variant that influences two different traits, trait A and trait B. In vertical pleiotropy, the traits themselves are biologically related, so that the variant’s effect on trait A actually causes the effect on trait B. A key feature of vertical pleiotropy is that two traits that are biologically related should be related regardless of which specific gene or variant is causing the effect. This induces correlation between GWAS effect sizes on the two traits across an entire set of variants. For example, we expect that *any* variant that increases LDL cholesterol also increases risk of coronary artery disease, because we suspect that it is the increase in LDL cholesterol itself that causes increased disease risk. This results in a correlation between variant effect sizes for LDL cholesterol and coronary artery disease, which has been detected in multiple studies (11–13). The methodology of Mendelian Randomization uses this predicted correlation within a given set of variants to formulate a statistical test for causal relationships among traits, which is now widely used for biological discovery (14,15). We extend this methodology to use the entire set of SNVs evaluated by GWAS, treating a GWAS-wide correlation between two traits as evidence of a vertical pleiotropic relationship between these traits.

**Figure 1:**
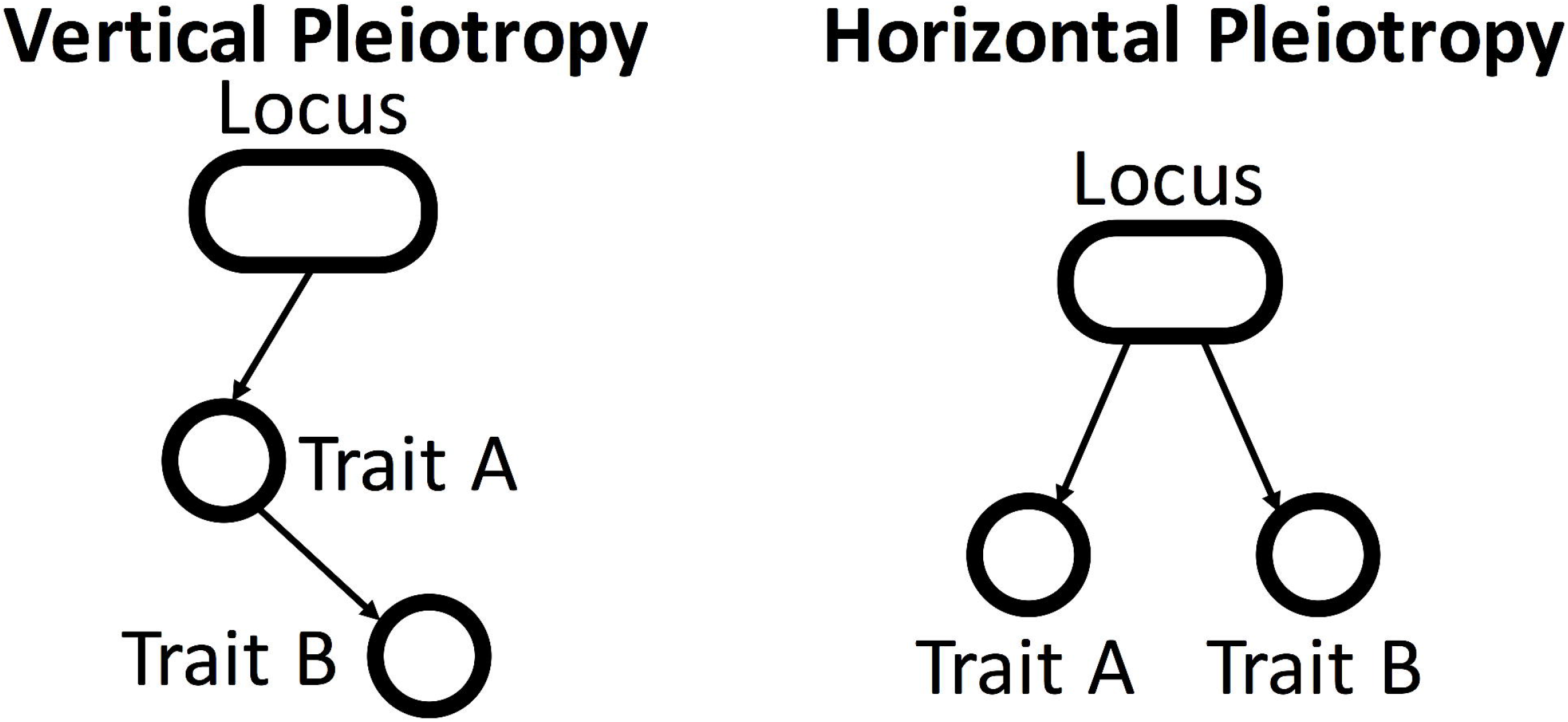
Schematic of different types of pleiotropy. Previous studies distinguish between vertical pleiotropy, where effects on one trait are mediated through effects on another trait, and horizontal pleiotropy, where effects on multiple traits are independent.

In the case of horizontal pleiotropy, an individual variant acts on traits A and B without mirroring any trait-level relationship between them. Unlike vertical pleiotropy, since we are not considering the variant-level effect as evidence of a relationship between the two traits, we cannot detect horizontal pleiotropy by detecting correlations between traits. Instead, each horizontally pleiotropic variant acts by its own unique mechanism. These particular pleiotropic variants, therefore, should show a relationship between the two traits that deviates from the relationship we would infer from the genome-wide correlation of effect sizes between them. This deviation from the correlation between traits is not a prediction of any kind of model of pleiotropy, but simply follows from our definition of the term “horizontal pleiotropy”: any pair of traits whose effect sizes are correlated across all variants is by definition related by vertical pleiotropy, while any variant whose effects on two traits substantially deviate from the trait-level relationship between those traits is by definition exhibiting horizontal pleiotropy.

### A quantitative score for pleiotropy

We have developed a method to measure horizontal pleiotropy using summary statistics data from GWASs on multiple traits. Our method relies on applying a statistical whitening procedure to a set of input variant-trait associations, which removes correlations between traits caused by vertical pleiotropy and normalizes effect sizes across all traits. Using the decorrelated association Z-scores, we measure two related but distinct components of pleiotropy: the total magnitude of effect on whitened traits (“magnitude” score, denoted *P_m_*) and the total number of whitened traits affected by a variant (“number of traits” score, denoted *P_n_*). Both scores are then scaled by the number of traits and multiplied by 100, so that the final score represents the value as it would be measured in a dataset of 100 traits. This two-component quantitative pleiotropy score allows us to measure both the magnitude (pleiotropy magnitude score *P_m_*) and quantity (pleiotropy number of traits score *P_n_*) of horizontal pleiotropy for all SNVs in the human genome. In principle these are distinct quantities: the magnitude score *P_m_* measures the total pleiotropic effect size of a variant across all traits, while the number of traits score *P_n_* measures the number of distinct pleiotropic effects a variant has. A variant with a high *P_m_* score and a low *P_n_* score has a large effect spread over a small number of traits; a variant with a low *P_m_* score and a high *P_n_* score has only a minor effect overall, but that effect is spread out across a large number of traits; and a variant with high scores on both components has a large effect that is spread across a large number of traits. Since we expect these scores to be heavily influenced by linkage disequilibrium (LD), we regress *P_m_* and *P_n_* against LD scores to produce an LD-corrected score (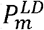 and 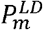) (**Figures 2, 3; Methods**).

**Figure 2:**
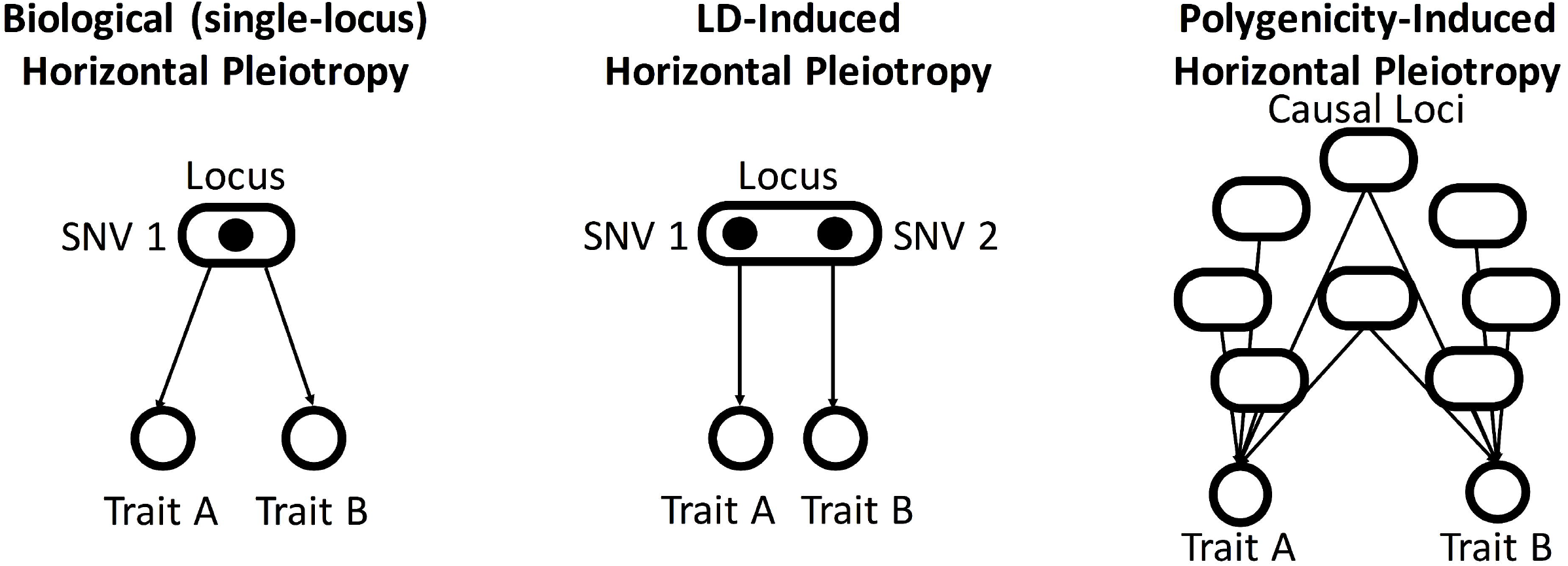
Contributions of linkage disequilibrium (LD) and polygenicity to horizontal pleiotropy. In addition to the normal sense of horizontal pleiotropy, both linkage disequilibrium (LD) and polygenicity are expected to contribute to horizontal pleiotropy. In the case of LD-induced horizontal pleiotropy, two linked SNVs have independent effects on different traits which appear pleiotropic because of the linkage between the SNVs. In the case of polygenicity-induced horizontal pleiotropy, two highly polygenic traits have an overlap in their polygenic footprint.

**Figure 3:**
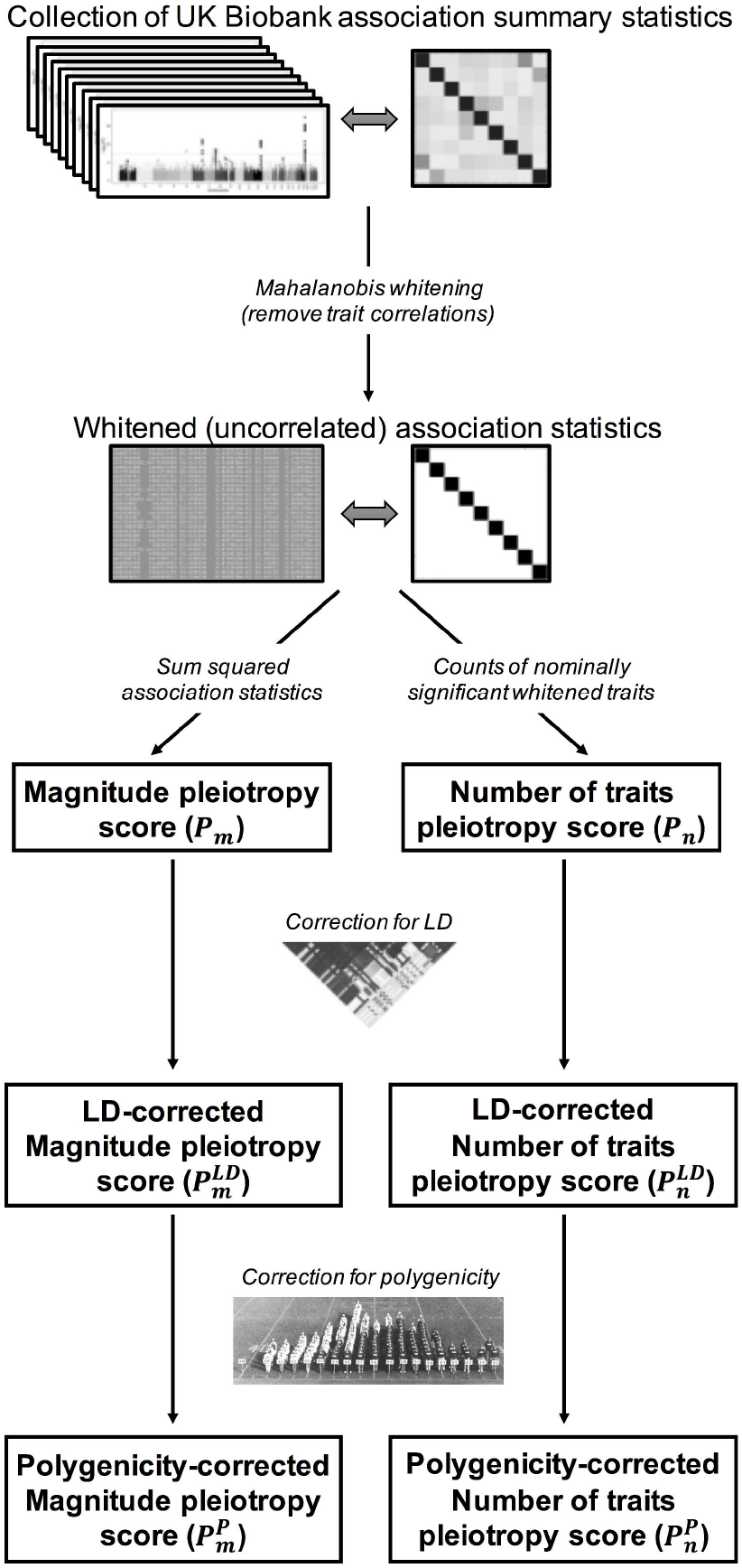
Two component pleiotropy score method. We (i) collect association statistics from the UK Biobank, (ii) process them using Mahalanobis whitening, (iii) compute the two components of our pleiotropy score (*P_m_* and *P_n_*) based on the whitened association statistics, (iv) use LD scores to correct for LD-induced pleiotropy (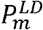 and 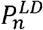), and (v) use permutation-based *P*-values to correct for polygenic architecture (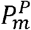 and 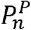).

### Calculating significance of pleiotropy

We compute *P*-values for the two components of our pleiotropy score using two different procedures, corresponding to two different null expectations.

1. Theoretical P-values (Raw pleiotropy score [*P_m_* and *P_n_*] or LD-corrected pleiotropy score [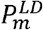 and 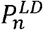]), calculated analogously to *P*-values for genetic association studies including GWAS, based on a null scenario where variants do not exhibit pleiotropic effects on observed traits.
2. Empirical *P*-values (Polygenicity/LD-corrected pleiotropy score [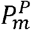 and 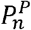), calculated by permutation of the observed distributions of whitened traits. These *P*-values are based on a null scenario where variants may have significant effects on one or more traits, but the effects of each variant on each trait are independent and the number of variants with effects on multiple traits is no more than would be expected by chance.

This empirical correction for polygenicity is required because polygenicity is a major factor that can produce pleiotropy. For example, it has been estimated that approximately 100,000 independent loci are causal for height in humans (16). If the total number of independent loci in the human genome is approximately 1 million, this corresponds to about 10% of the human genome having an effect on height. If we imagine multiple phenotypes with this same highly polygenic genetic architecture, we should expect substantial overlap between causal loci for multiple different traits, even in the absence of any true causal relationship between the traits, resulting in horizontal pleiotropy (**Figure 2**).

### Power to detect pleiotropy in simulations

We conducted a simulation study to evaluate the performance of our two-component pleiotropy score. We simulated 800,000 variants controlling 100 traits, varying the per-trait liability scale heritability of all traits *h*^2^ and the proportion of pleiotropic and non-pleiotropic causal variants. To introduce LD in the simulations, we used real LD architecture from 800000 SNVs from 1000 Genomes European population. We simulated Z-scores independently for each SNV and then propagate LD for a given SNV by “contaminating” its Z-score according to the Z-scores of the SNVs in LD with it. Under the null model, all trait-variant associations were independent, and no horizontal pleiotropy was added. Under the added-pleiotropy models, we randomly chose a fraction of causal variants and forced them to have simultaneous associations with multiple traits. The simulation study showed that both components of the pleiotropy score were well-powered to detect horizontal pleiotropy (**Figure 4**), and that the LD correction dramatically reduces the dependence of the pleiotropy score on LD (**Additional File 1: Fig. S1**). Under the null hypothesis of no added horizontal pleiotropy, the false positive rate was well controlled for both scores when there was low heritability or few causal variants. However, when there are many causal variants and high per-variant heritability, the LD-corrected pleiotropy score (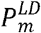 and 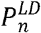) detects a large excess of pleiotropic variants, due to serendipitous overlap between causal variants without explicitly induced pleiotropy. The LD/polygenicity-corrected empirical *P*-value (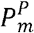 and 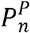) does not detect this serendipitous pleiotropy at the same high rate.

**Figure 4:**
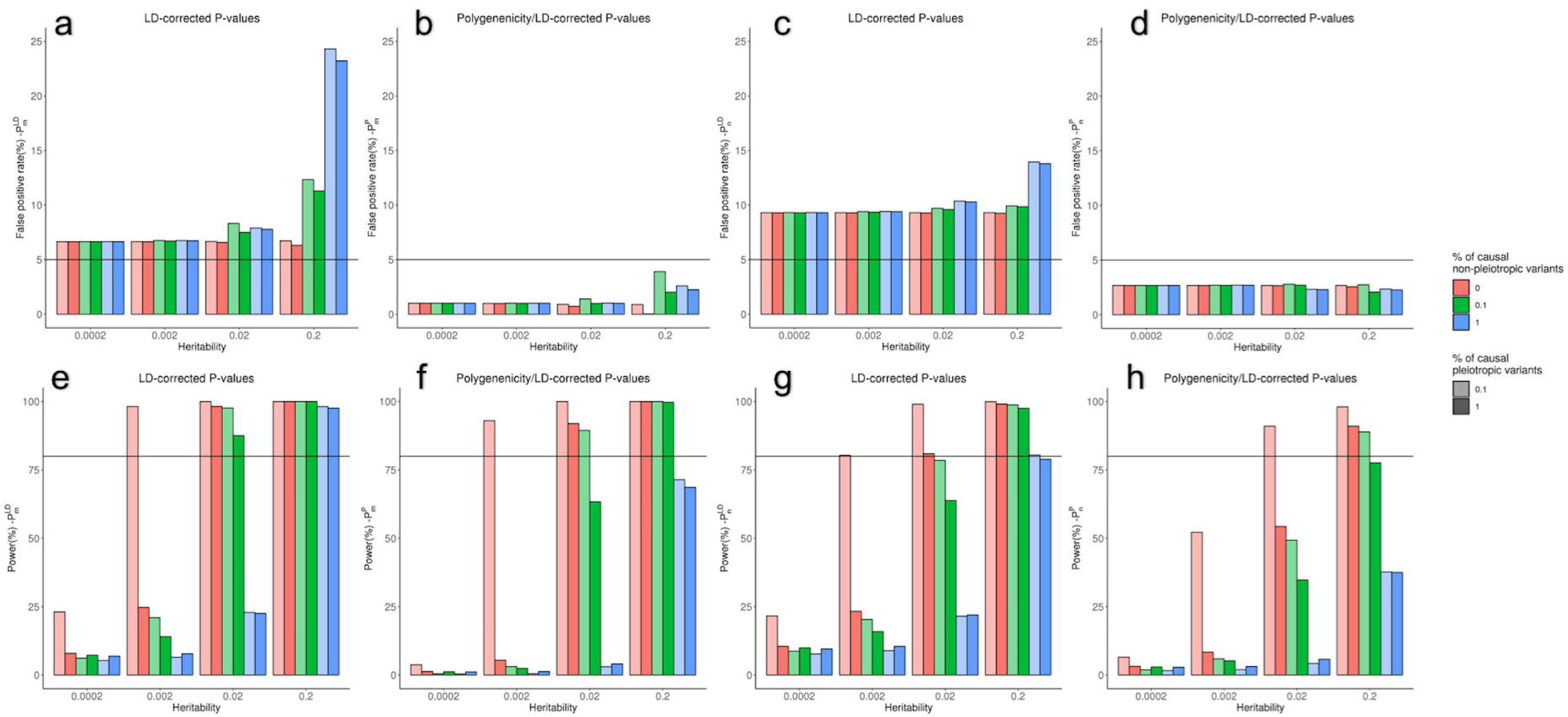
Simulation study showing false positive rate (a,b,c,d) and power (e,f,g,h) of two-component pleiotropy score. Top row shows performance on non-pleiotropic simulated variants (black line shows 5% false positive rate); bottom row shows performance on pleiotropic variants (black line shows 80% power). Simulations were run for both 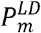 (left) and 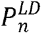 (right), and both without correction for polygenicity (a,c,e,g) and with the correction (b,f,d,h), with per-variant heritability ranging from 0.0002 to 0.2, proportion of non-pleiotropic causal loci ranging from 0 to 1%, and proportion of pleiotropic causal loci ranging from 0.1% to 1%. Our method has good power to detect pleiotropy for highly heritable traits, though its power is reduced by extreme polygenicity. Extreme polygenicity also increases the false positive rate, though this effect is corrected by our polygenicity correction.

In the presence of added horizontal pleiotropy, our approach was powered to detect pleiotropy with per-variant heritability *h*^2^ as small as 0.002 if there are no non-pleiotropic causal variants. In the presence of both pleiotropic and non-pleiotropic causal variants, detecting pleiotropy was more difficult, but our approach still had appreciable power to detect pleiotropic variants, which increased with increasing per-variant heritability and decreased with increasing numbers of non-pleiotropic causal variants. Adding the correction for polygenic architecture (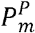 and 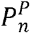) reduced this power only slightly. The power of our method was not substantially reduced by increasing the number of traits affected by pleiotropic variants (**Additional File 1: Fig. S2**) or by adding a realistic correlation structure between the traits (**Additional File 1: Fig. S3**).

### Genome-Wide Pleiotropy Study (GWPS) reveals pervasive pleiotropy

To apply our method to real human association data, we used GWAS association statistics for 372 heritable medical traits measured in 337,119 individuals from the UK Biobank (17–19). We successfully computed our two-component pleiotropy score for 767,057 variants genome-wide and conducted a genome-wide pleiotropy study (GWPS), by analogy to a standard GWAS (**Figure 3; Methods**). **Additional File 1: Fig. S4** shows the resulting quantile-quantile plots (Q-Q plots). We observed significant inflation for both the LD-corrected magnitude score 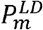 and number of traits score 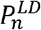 (Mann-Whitney U test *P* < 10^-300^ for both). Furthermore, we observed across both scores that horizontal pleiotropy was widely distributed across the genome, rather than being localized to a few specific loci (**Additional File 1: Fig. S5**). Testing an alternative strategy for computing the phenotype-correlation matrix using all SNVs produced comparable results (Pearson *r* = 0.995 and 0.964 for 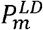 and 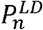 respectively) to our strategy of using a pruned set of SNVs to account for LD (*r*^2^ < 0.1) (**Additional File 1: Fig. S6**).

### Pleiotropy is driven by polygenicity

We applied the permutation-based empirical P-value calculation (Polygenicity/LD-corrected pleiotropy score: 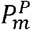 and 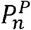) to correct for the known polygenic architecture of traits and test whether any loci are pleiotropic to a greater extent than would be expected due to polygenicity. **Additional File 1: Figs. S7 and S8** show the resulting Q-Q plots and Manhattan plots. In contrast to the results from the LD-corrected pleiotropy score (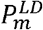 and 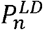), we do not find pleiotropy significantly in excess of what would be expected from the known polygenic architecture of traits: there are dramatically fewer loci with genome-wide significant levels of pleiotropy after correcting for polygenic architecture, and the genome-wide distribution of pleiotropy score shows less pleiotropy than expected (Mann-Whitney U test *P* < 10^-300^ for both 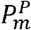 and 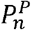).

As an additional test of whether the pleiotropy we observe is driven by polygenicity, we calculated the polygenicity of the same 372 heritable traits from the UK Biobank. We measured polygenicity using a version of the genomic inflation factor corrected using LD score 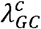 (20). We then stratified these traits by 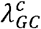 after controlling for heritability (Methods), and calculated the two-component LD-corrected pleiotropy score [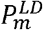 and 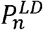]) and *P*-values for each component independently for every variant in the genome using each of these bins of traits. We observed that both scores are highly dependent on polygenicity, with the lowest-polygenicity bins in each heritability class showing very little inflation. (**Figure 5; Additional File 1: Table S1**). Taken together, these results suggest that extreme polygenicity drives horizontal pleiotropy, and that this has an extremely large effect on the genetic architecture of human phenotypes.

### Genome-wide distribution of pleiotropy score gives insight into genetic architecture

In addition to observing genome-wide inflation of the pleiotropy score, we can also gain insight from the distribution of the pleiotropy score on a more granular level.

**Figure 6a** shows the distribution of pleiotropy score for independent SNVs (LD pruned to a threshold of *r*^2^ < 0.1) compared to the expectation under the null hypothesis of no pleiotropic effect. We observe a large excess in the number of traits score 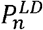, and a smaller but still highly significant excess in total magnitude of pleiotropic effect 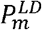. This excess comes in part from a long tail of highly pleiotropic loci that pass the threshold of genome-wide significance (dashed line in **Figure 6a**), but is primarily driven by weak pleiotropy among loci that do not reach genome-wide significance.

**Figure 5:**
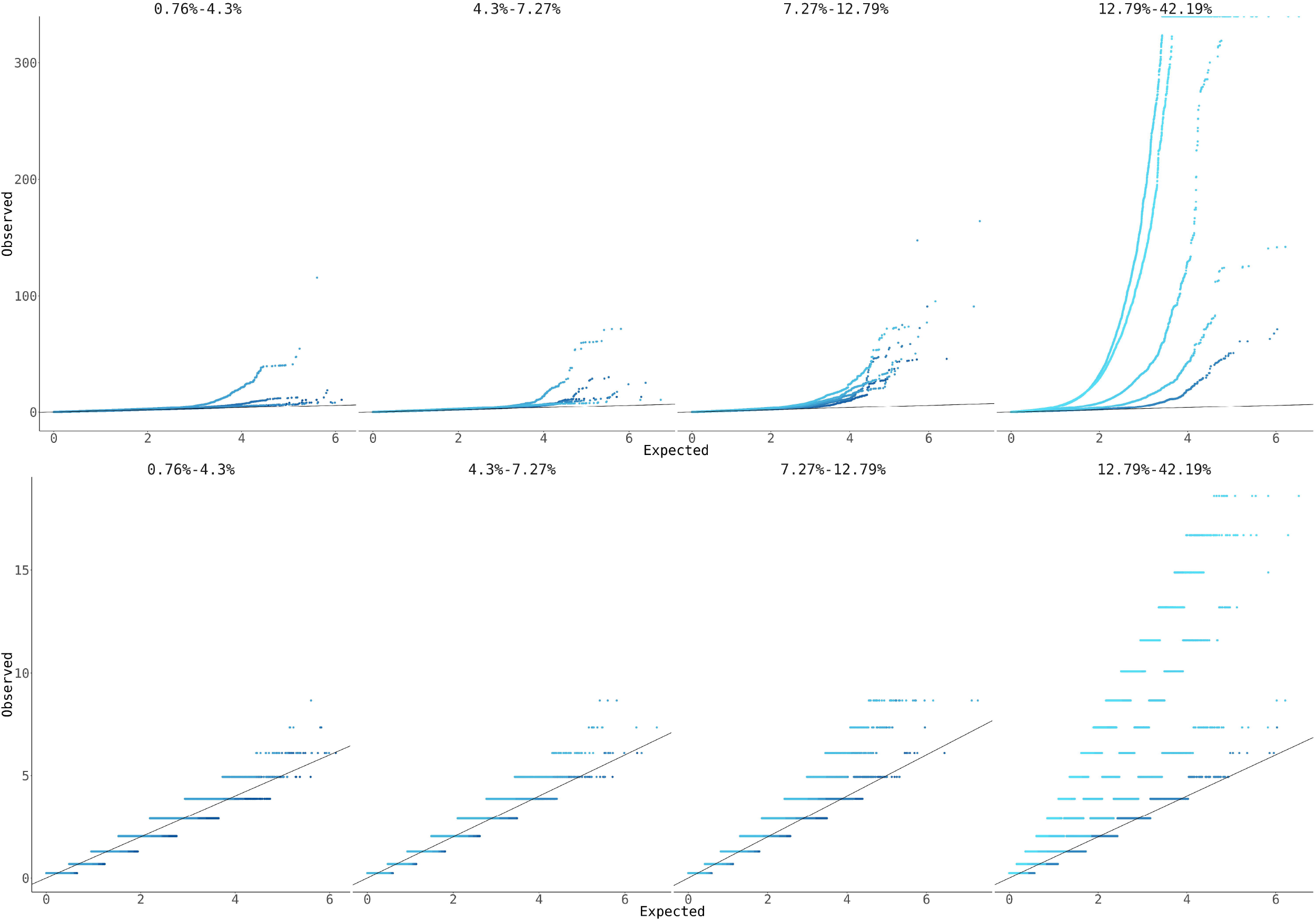
Quantile-quantile (Q-Q) plots showing the inflation of the pleiotropy score as a function of polygenicity. Variants are stratified into 4 batches of about 80 traits each by heritability, and then subdivided into 5 batches of about 20 traits each by polygenicity, as measured by corrected genomic inflation factor 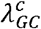. Darker shades represent low polygenicity and lighter shades represent high polygenicity. All panels show -log_10_ transformed *P*-values. The black lines show the expected value under the null hypothesis.

### Pleiotropy score is correlated with molecular and biological function

To further investigate the properties of pleiotropic variants, we examined the effects of various functional and biochemical annotations on our LD-corrected pleiotropy score (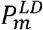 and 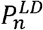) (**Table 1; Methods**). Using annotations from Ensembl Variant Effect Predictor (21), we observed that both components of the pleiotropy score are higher on average in transcribed regions (coding and UTR) than in intergenic noncoding regions. This result was confirmed and expanded by annotations from Roadmap Epigenomics (22), which showed that regions whose chromatin configurations were associated with actively transcribed regions, promoters, enhancers, and transcription factor binding sites had significantly higher levels of both components of the pleiotropy score, while heterochromatin and quiescent chromatin states had significantly lower levels. Investigating individual histone marks, we found that both the repressive histone mark H3K27me3 and the activating histone mark H3K27ac were associated with elevated levels of pleiotropy, although the activating mark H3K27ac had a larger effect. This may indicate that being under active regulation at all produces higher levels of pleiotropy, whether that regulation is repressive or activating.

We also used data from the Genotype-Tissue Expression (23) project to measure the connection between transcriptional effects and our pleiotropy score (**Table 1**). Consistent with the previous observation that functional regions had higher pleiotropy scores, we found that variants that were identified as *cis*-eQTLs for any gene in any tissue had higher pleiotropy scores on average. Within eQTLs, we also observed significant correlations between our pleiotropy score and the numbers of genes (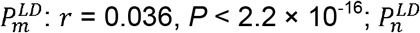: *r* = 0.035, *P* < 2.2 × 10^-16^) and tissues (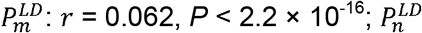: *r* = 0.059, *P* < 2.2 × 10^-16^) where the variant was annotated as an eQTL, showing that our pleiotropy score is related to transcriptional measures of pleiotropy.

**Table 1:**
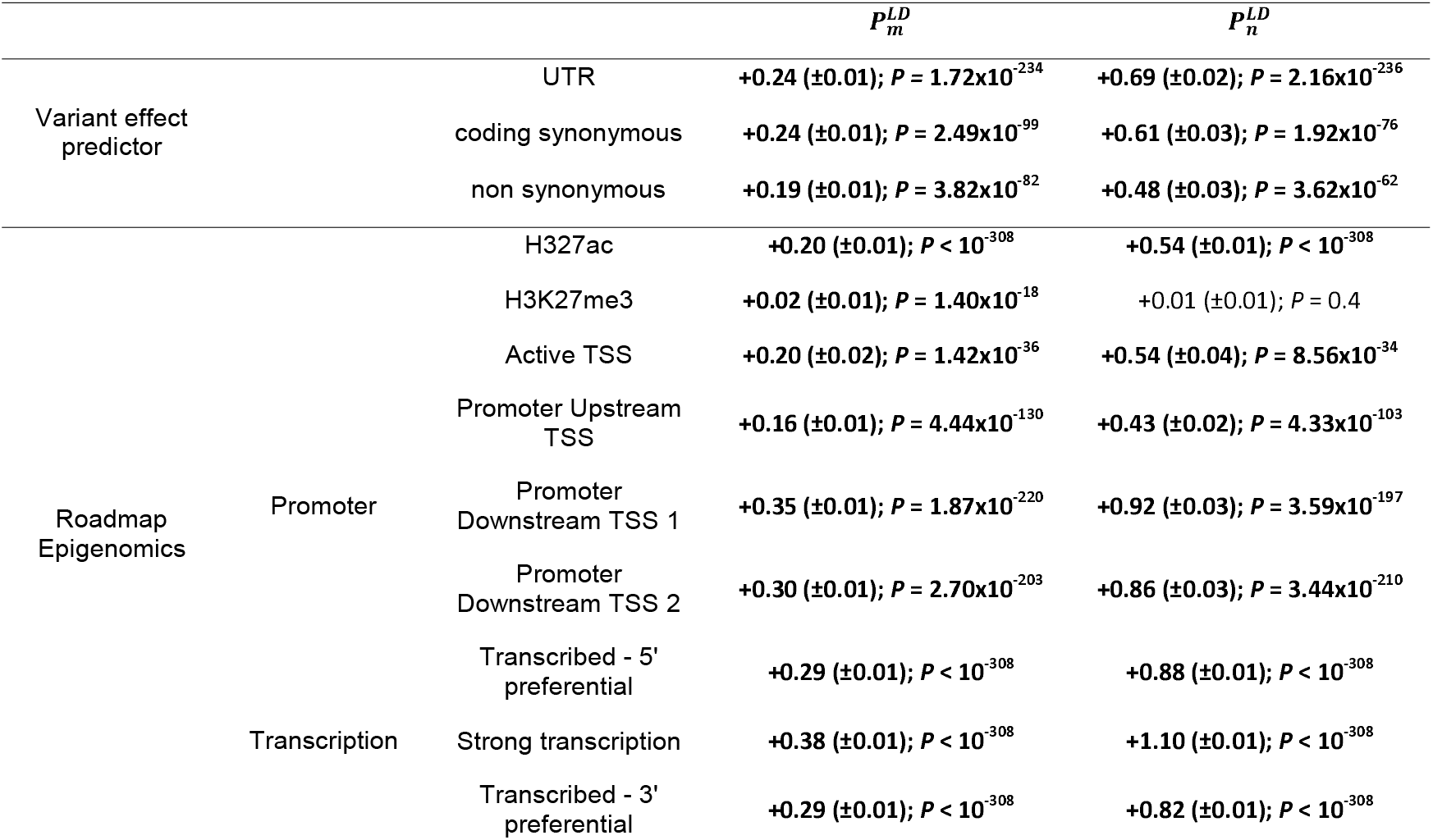
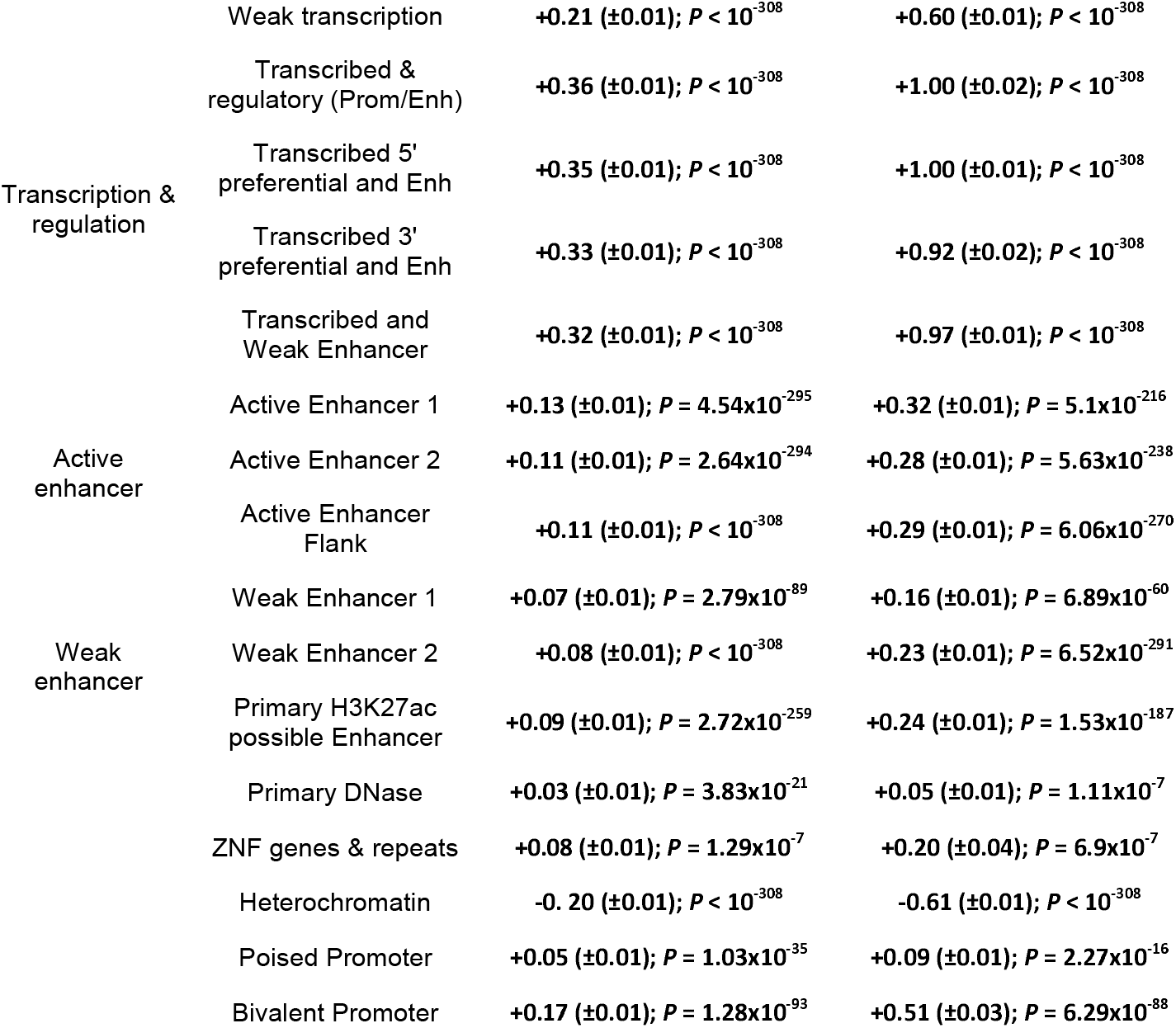
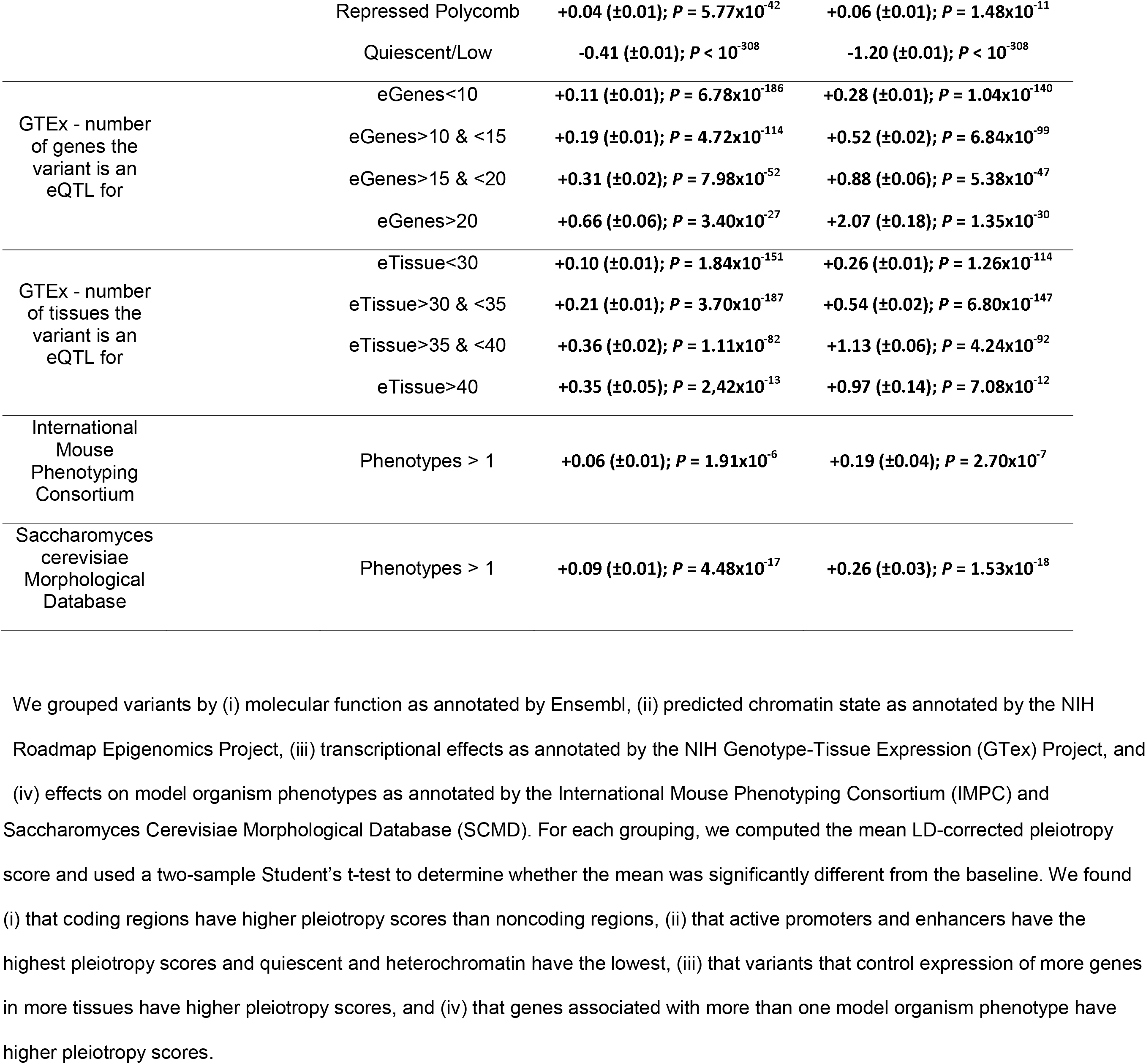
Functional enrichment analysis of pleiotropy score.

Finally, we found that variants that are eQTLs for genes whose orthologs are associated with multiple measurable phenotypes in mice or yeast have higher pleiotropy scores, demonstrating that our pleiotropy score is also related to pleiotropy in model organisms.

All these results are consistent when using the Polygenicity/LD-corrected pleiotropy score (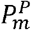 and 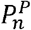), indicating that the association of pleiotropy with molecular and biological function is not exclusively driven by highly polygenic architecture (**Additional File 2**).

### Genome-wide pleiotropy study identifies novel biological loci

By analogy to standard GWAS, our GWPS methodology can identify individual variants that have a genome-wide significant level of horizontal pleiotropy. Using the LD-corrected magnitude score 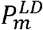, we identified 74,335 variants in 8,093 independent loci with a genome-wide significant level of horizontal pleiotropy, while using the LD-corrected number of traits score 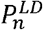 identified 18,393 variants in 2,859 independent loci with a genome-wide significant level of horizontal pleiotropy, all of which are also identified by the LD-corrected magnitude score 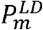 (**Methods, Additional File 1: Table S2**). Applying the same analysis to the Polygenicity/LD-corrected pleiotropy score, using the Polygenicity/LD-corrected magnitude score 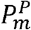 identified no genome-wide significant loci, but using the Polygenicity/LD-corrected number of traits score 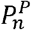 identified 2,674 variants in 432 loci. Strikingly, a majority of loci significant in 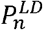 (1,519 of 2,859) or 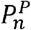 (294 of 432), along with a sizeable minority of loci significant in 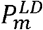 (2,934 of 8,093), have no entry in the NHGRI-EBI GWAS catalog, meaning that they have never been reported as an associated locus in any published GWAS. These loci represent an under-recognized class of genetic variation that has multiple weak to intermediate effects that are collectively significant, but no specific strong effect on any one particular trait. Functional enrichment analysis on genes near these genome-wide significant loci implicates a wide range of biological functions, including cell adhesion, post-translational modification of proteins, cytoskeleton, transcription factors, and intracellular signaling cascades (**Additional File 3**). Loci significant in 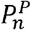 show a more focused subset of functions, with a greater role for nuclear proteins regulating transcription and chromatin state, suggesting that these are the functions that exhibit horizontal pleiotropy beyond the baseline level induced by polygenicity. The role of these novel loci and these biological processes in human genetics and biology may be a fruitful area for future study, with the potential for biological discovery.

### Pleiotropic loci replicate in independent GWAS datasets

As replication datasets, we used two additional sources of GWAS summary statistics to calculate our LD-corrected pleiotropy score (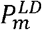 and 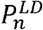): previously published GWASs and meta-analyses for 73 human complex traits and diseases, which we collected and curated manually from the literature (**Methods, Additional File 1: Table S3**) (24); and a previously published study of 430 blood metabolites measured in 7,824 European adults (25). For all variants covered by the UK Biobank, we were able to compute our pleiotropy score independently using these two datasets (**Figure 7**). In the traits and diseases dataset, we observed that 57% of 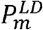 loci and 38% of 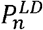 loci replicated, while in the blood metabolites dataset, we observed that 17% of 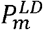 loci and 12% of 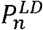 loci replicated, compared to 5% of 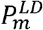 loci and 6% of 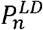 loci expected by chance according to a permutation-based null model. This high level of replication using independent sets of GWAS summary statistics suggests that our pleiotropy score is capturing an underlying biological property, rather than an artifact of the UK Biobank study.

**Figure 7:**
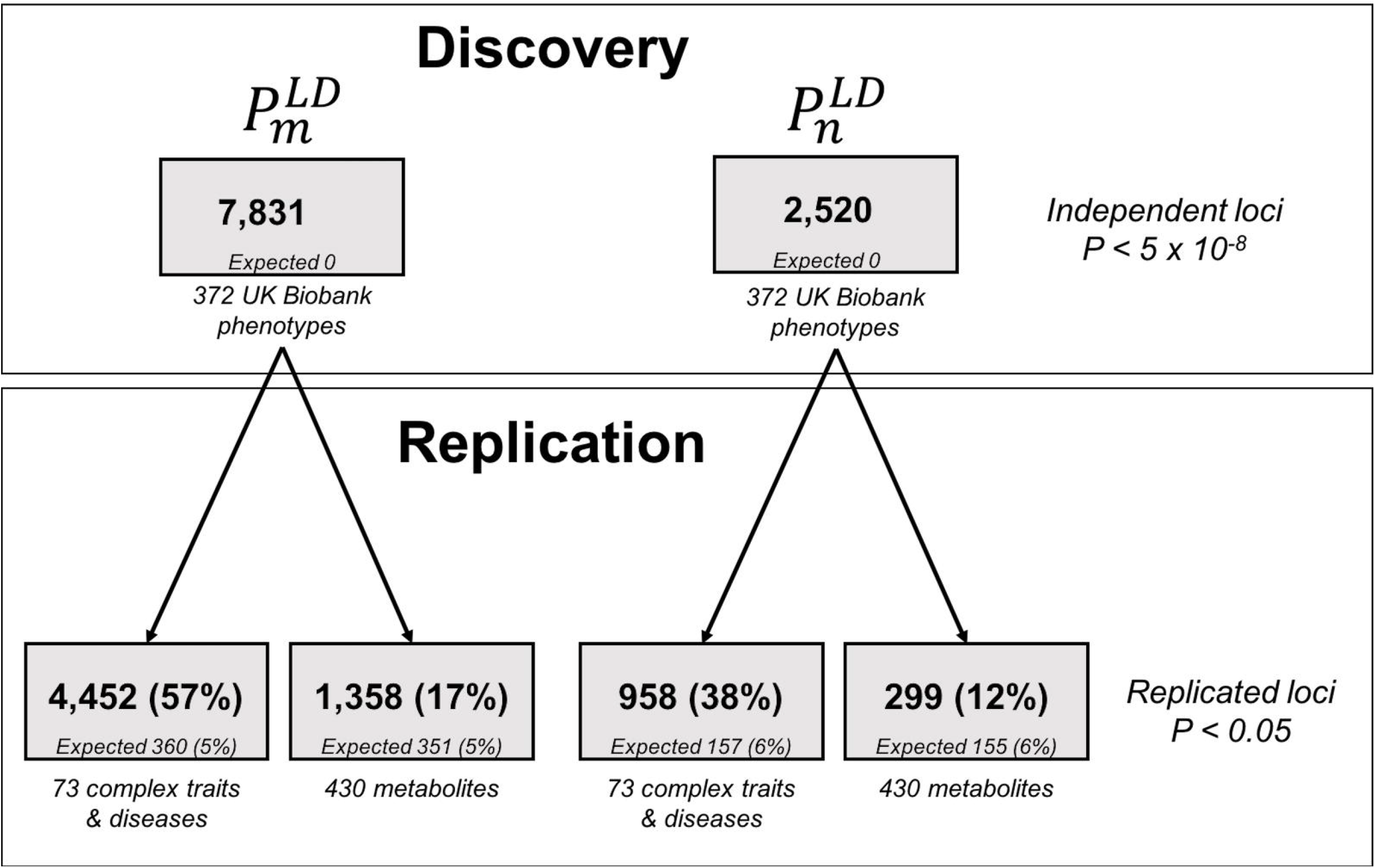
Replication analysis for the genome-wide pleiotropy study. We used 372 UK Biobank heritable medical traits as our discovery dataset, and independent datasets of 73 complex traits and diseases and 430 blood metabolites as replication datasets. In each case, expected fraction of replication was empirically determined using a permutation analysis.

### Pleiotropy is correlated with specific complex traits and diseases

To characterize the phenotypic associations of these loci, we used our replication dataset of published GWAS summary statistics for 73 human quantitative traits and diseases, plus nine additional traits we excluded from our replication dataset for a total of 82 (**Methods**). We are not able to compute directly the degree of pleiotropy exhibited by these traits, since our definition of horizontal pleiotropy applies only to individual variants and does not apply to traits. However, we can identify traits whose GWAS variant associations are correlated to our pleiotropy score, which in some sense represents the traits that contribute most to our signal of pervasive horizontal pleiotropy. **Figure 6c** shows the correlations between our LD-corrected pleiotropy score (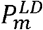 and 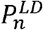) and the association statistics for these 82 traits and diseases. The most strongly correlated traits were anthropometric traits like body mass index, waist and hip circumference, and height; certain blood lipid levels, including total cholesterol and triglycerides; and schizophrenia. These are all known to be highly polygenic and heterogeneous traits. The least correlated traits include several measurements of insulin sensitivity and glucose response, such as the insulin sensitivity index (ISI), certain features of brain morphology, and the inflammatory biomarker lipoprotein(a). This may be partly due to low sample size of the corresponding GWASs. However, these correlations do not appear to be driven exclusively by sample size: in cases where multiple GWASs for the same trait have been performed on subsamples of the population (for example, males only, female only, and combined), the sample size only marginally affects the correlation (**Additional File 1: Table S4**). Another contributing factor may be heritability: height, in particular, is among the most heritable traits we examined, while ISI and the brain morphology features are among the least.

**Figure 6:**
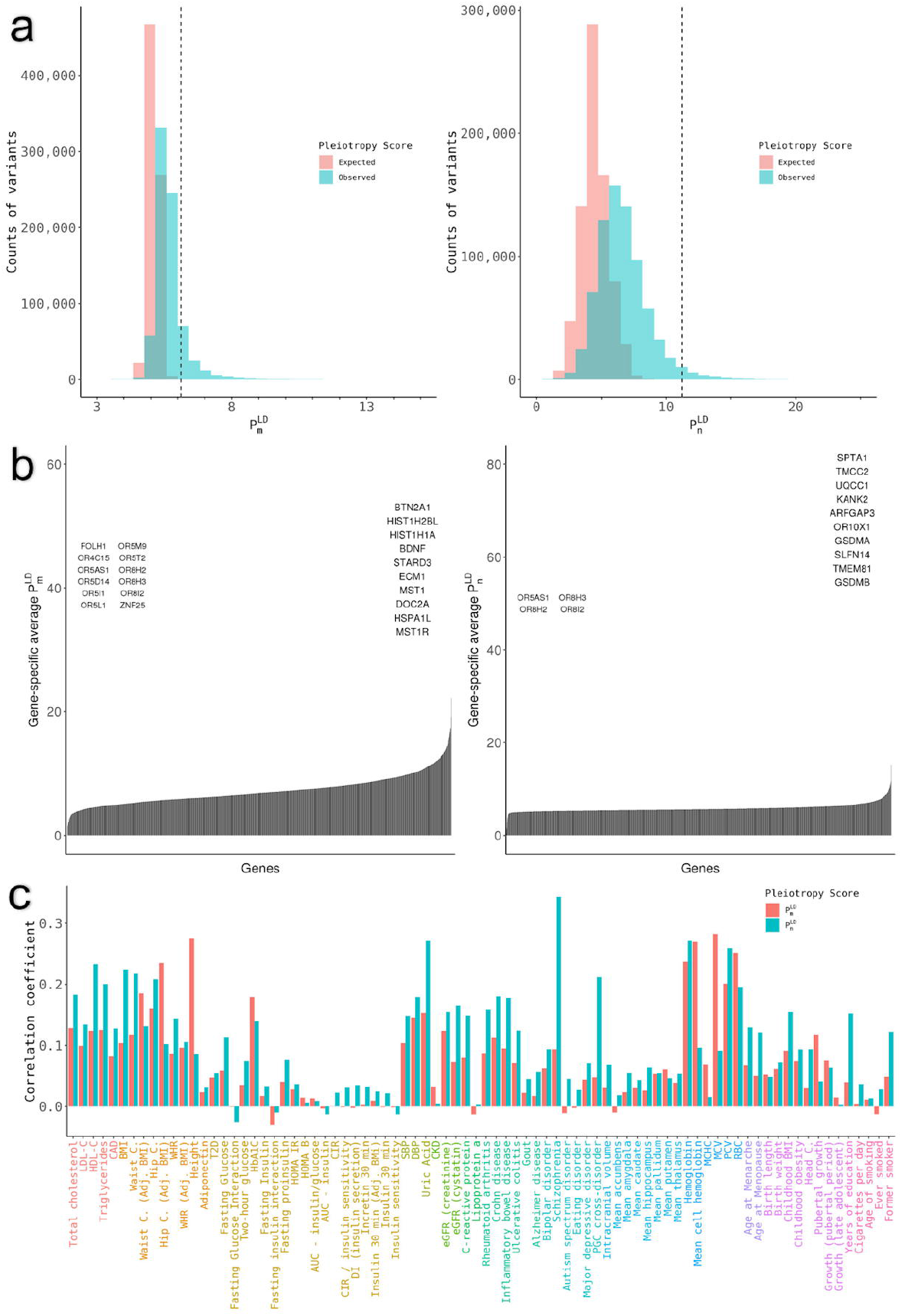
Distribution of the pleiotropy score among variants (a), genes (b), and traits (c). Panel a shows the global distribution of 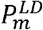 (left) and 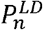 (right) for the 767,057 tested variants. The expected distribution under the null hypothesis of no pleiotropy is shown in red and the observed distribution is shown in blue. The vertical line represents the value of the pleiotropy score corresponding to genome-wide significance (*P* < 5 × 10^-8^). 1,769 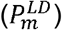 and 643 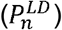 variants are not represented for the sake of clarity, because they have extreme values for the pleiotropy score. Panel b shows the distribution of the average pleiotropy score for coding variants in each gene for 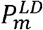 (left) and 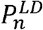 (right). The top ten genes are represented on the right side of the plots, whereas genes with a pleiotropy score of 0 are represented on the left side of the plots. Panel c shows the contribution of pleiotropic variants to 82 complex traits and diseases. Contribution of pleiotropic variants is calculated as the correlation coefficient between the absolute value of Z-scores and the pleiotropy score among variants that are genome-wide significant for the pleiotropy score (*P* < 5 × 10^-8^ for 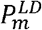 and 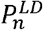 respectively).

## Discussion

We have presented HOPS, a framework for scoring horizontal pleiotropy across human genetic variation. In contrast to previous analyses, our framework explicitly distinguishes between horizontal pleiotropy and vertical pleiotropy or biological causation. After applying HOPS to 372 heritable medical traits from the UK Biobank, we made the following observations: 1) horizontal pleiotropy is pervasive and widely distributed across the genome; 2)) horizontal pleiotropy is driven by extreme polygenicity of traits; 3) horizontal pleiotropy is significantly enriched in actively transcribed regions and active regulatory regions, and is correlated with the number of genes and tissues for which the variant is an eQTL; 4) there are thousands of loci that exhibit extreme levels of horizontal pleiotropy, a majority of which have no previously reported associations; and 5) pleiotropic loci are enriched in specific complex traits including body mass index, height, and schizophrenia. These findings are largely consistent between the magnitude of pleiotropy score *P_m_* and the number of traits score *P_n_*, although we note some differences where some variants are primarily associated with 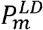 but not 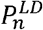. This indicates that these signals are driven by loci that both influence a large number of traits and have relatively large combined effects, and secondarily by loci that have large combined effects but only influence a handful of traits each, with minimal contribution from loci that influence a large number of traits but have small combined effects. Conversely, after applying the correction for polygenicity, we only observe variants that are significant for 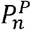, but not for 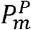. This indicates that, while there do exist horizontal pleiotropic master control loci that affect more traits than we would expect from the random overlap of multiple highly polygenic traits, the overall effect of these loci is not noticeably larger than we would expect.

This analysis is enabled by the technique of whitening trait associations to remove correlations between traits. This lets us count pleiotropic effects in a more objective and systematic way, as opposed to manually selecting putatively independent traits to count, or manually grouping traits into independent blocks. However, it does come with three major limitations compared to these approaches. First, it is somewhat more difficult to tell which specific traits are driving a signal of pleiotropy at a particular locus. Our whitened traits are combinations of real observed traits, and do not necessarily correspond to any specific biological traits of interest. However, it is relatively easy to inspect the input GWAS summary statistics for a particular variant of interest to see which traits it is associated with. Furthermore, since pleiotropic loci are by definition associated with a large cross-section of traits, this kind of inspection is not likely to be very informative about specific traits. Second, the whitening procedure has the counterintuitive property that a variant that has a narrow effect on a single trait without also affecting correlated traits can appear to be highly pleiotropic. For example, if a variant had a strong risk-increasing effect on coronary artery disease (CAD), but no effect on any of the known upstream risk factors of CAD (such as blood lipid levels or adiposity) or any of the known downstream consequences of CAD (such as inflammatory biomarkers or increased mortality), such a variant would appear as highly pleiotropic in our analysis. Our analysis would interpret the variant as increasing the risk of CAD while suppressing these upstream and downstream factors. We believe this treatment is appropriate, however counterintuitive. Regardless, these kinds of isolated effects are fairly rare: in our dataset of 372 heritable traits from UK Biobank, only 6% of variants (42,684 of 767,057) reach genome-wide significance for only a single trait. Indeed, it is unlikely by definition that a variant is associated with only one trait from a set of correlated traits, since we compute our correlations from observed association statistics. Third, we assume all genetic effects are additive and independent, and we do not model epistasis or other more complex genetic architectures.

Our findings are in keeping with several recent studies that have found abundant pleiotropy in the genome (26,27,8,2,9). HOPS goes a step further than many of these studies by explicitly removing vertical pleiotropy between traits, which are indicative of fundamental biological relationships between traits (8,24,28). Furthermore, the current study has evaluated horizontal pleiotropy in human genetic variation genome-wide, whereas previous studies have focused on only a small subset of disease-associated variants identified from GWAS. Our results therefore suggest that there is substantial complexity and heterogeneity not only in causal relationships between human traits, but also in the genetic architecture of individual traits.

Our findings have several important implications for the field of human genetics. First, our observation of ubiquitous horizontal pleiotropy is problematic for Mendelian Randomization (MR) methods, which assumes horizontal pleiotropy to be absent. Recent developments in the field of MR include methods that account for horizontal pleiotropy explicitly (24,28,29); our results reinforce the importance of these methods. The presence of widespread horizontal pleiotropy suggests that single-instrument methods that independently account for every variant, each of which presumably has pleiotropic effects on many different distinct traits, should be considered in addition to multi-instrument methods for MR, which collapse many variants into a single polygenic score for analysis, and therefore treat all variants equivalently.

Second, our results appear to support the “network pleiotropy” hypothesis of Boyle, Li, and Pritchard (16), which proposes widespread pleiotropy driven by small perturbations of densely connected functional networks, where any perturbation in a relevant cell type will have at least a small effect on all phenotypes affected by that cell type. A subsequent paper detailed a more specific mechanism, where causal effects are driven by many biological components that are only indirectly related to the phenotype itself (30). Many of the functional enrichments we observe, including transcription factors, cytoskeleton, and intracellular signaling cascades, represent components that can plausibly influence a wide variety of cell types and processes, providing evidence for this model over one where a specific biological component is largely responsible for pleiotropy. The fact that the magnitude of pleiotropy score *P_m_* and the number of traits score *P_n_* give largely consistent results also supports this model, where a larger biological effect in a given tissue will perturb a greater number of phenotypes relevant to that tissue, although we note that some variants have high magnitude of pleiotropy score *P_m_* and low number of traits score *P_n_*, which may represent a small class of variants that has large biological effects without perturbing a large number of phenotypes.

While our results largely support this network pleiotropy hypothesis, we have also demonstrated an alternate view of horizontal pleiotropy in the context of highly polygenic causation. In our simulations, introducing extreme polygenicity at the levels suggested by these papers inherently results in high levels of horizontal pleiotropy detectable by our score, independent of any assumptions about the mechanism of pleiotropy or of polygenicity. Indeed, our null hypothesis of no horizontal pleiotropy, that 5% of the genome is independently causal to each trait with *P* < 0.05, is trivially rejected when a single trait is influenced by an unexpectedly large fraction of the genome. This means that, on some level, widespread horizontal pleiotropy in human genetic variation is simply a logical consequence of widespread polygenicity of human traits, regardless of the specific mechanism of either. In simple terms, the more loci are associated with each trait, the more chances there are for associations with multiple traits to overlap. Supporting this result, we find that controlling for the polygenic architecture of the input traits significantly attenuates our signal of pleiotropy, as does restricting to oligogenic traits. It may be the case that horizontal pleiotropy is only truly widespread among the most complex and polygenic subset of human traits.

## Conclusions

In this study, we have presented HOPS, a quantitative score for horizontal pleiotropy in human genome variation. Using this score, we have identified a genome-wide trend of highly inflated levels of horizontal pleiotropy, an underappreciated relationship of between horizontal pleiotropy with polygenicity and functional biology, and a large number of specific novel loci with high levels of horizontal pleiotropy. We expect further investigations using HOPS to yield deep insights into the genetic architecture of human traits and to uncover important novel biology.

## Methods

We developed a statistical method to measure horizontal pleiotropy using a two-component pleiotropy score. For a given variant, we measured 1) the total magnitude of pleiotropic effect the variant has and 2) the number of whitened traits affected by the variant.

### Z-scores decorrelation strategy

Observable traits and diseases can be highly correlated, which can lead to inflation of our pleiotropy score if the correlation is not properly accounted for. Therefore, we developed an efficient strategy to remove this correlation and obtain decorrelated traits. Let *Z^raw^* denote the matrix of raw Z-scores, with variants in columns and traits in rows, and *Σ* denote the corresponding correlation matrix between the Z-scores. Under the null hypothesis of no horizontal pleiotropy, Z-scores for each trait are assumed to follow a Gaussian distribution *N*(0,1), and the columns of *Z^raw^* collectively follow a multivariate Gaussian distribution *N*(0,*Σ*). Our goal is to eliminate the extra-diagonal terms of the correlation matrix *Σ*. To achieve this, we use a Mahalanobis whitening transformation on the matrix *Z^raw^* to obtain a whitened Z-score matrix *Z*. The procedure to obtain *Z* can be formally expressed as:

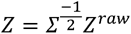

Under the null hypothesis of no horizontal pleiotropy, we expect *Z* to follow a multivariate Gaussian distribution *N*(0,*Id_l_*), where *Id_l_* is the identity matrix of size *l, l* being the number of traits.

In reality, the true correlation matrix *Σ* is unknown, and we must use an estimated correlation matrix 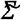 obtained by measuring the genome-wide correlation between actual Z-scores. We tested two approaches to obtain 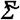, either using all genotyped variants genome-wide or using a subset of variants pruned to *r*^2^ < 0.1 in the 1000 Genomes European population to account for the effects of linkage disequilibrium (LD). Both approaches produced similar results (See **Additional File 1: Fig. S6**). In all subsequent analysis, we used covariance matrices estimated from pruned variants.

### Computation of the pleiotropy score

We computed two different scores to capture both the magnitude and number of traits of pleiotropy. First, we quantify the total pleiotropic magnitude of effect a variant using the magnitude pleiotropy score *p_m_*:

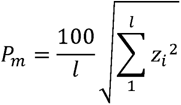

where *z_i_* is the whitened Z-score for trait *i* for a given variant. Second, we quantify the number of whitened traits affected by a variant using the number of pleiotropic traits score *P_n_*:

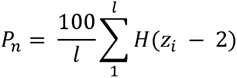

where *z_i_* is the whitened Z-score for trait *i* for the tested variant and *H*() is the Heaviside step function which equals 1 if |*z_i_*| > 2 and 0 otherwise. 2 represents a standard value of the Z-score which represents the normal threshold for nominal significance (*P* < 0.05).

### LD-corrected pleiotropy score

Similarly to LD score regression, each component of the pleiotropy score was regressed on the LD scores for all variants. Then, we regressed out the effect of LD on each component of the pleiotropy score independently to obtain an LD-corrected pleiotropy score. The LD-corrected pleiotropy score components 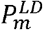 and 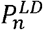 are given by:

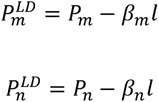

where *l* is the LD score of the variant site, and *β_m_* and *β_n_* are the regression coefficients for LD score on *P_m_* and *P_n_*, respectively.

### Computation of theoretical *P*-values for the pleiotropy score

Based on the observation that *Z* follows a multivariate standard Gaussian distribution *N*(0, *Id_l_*) under the null hypothesis of no pleiotropy, *P*-values can easily be computed for *P_m_* and *P_n_*. Under the null hypothesis, the square of *P_m_* (or 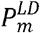) follows a chi-square distribution *χ*^2^(*l*) where *l* is the total number of traits. Likewise, *P_n_* (or *Pf*) follows a binomial distribution *B*(*l,p*) where *l* is the total number of traits and *p* the probability to get a Z-score greater than 2 under the standard Gaussian distribution (*P* ≈ 0.045).

### Computation of empirical (polygenicity/LD-corrected) *P*-values for the pleiotropy score

To correct for the known polygenic architecture of traits in addition to LD, we additionally computed empirical permutation-based *P*-values for both 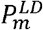 and 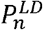. We performed 25 random permutations of the input Z-scores for each observable trait, producing millions of permuted variants. We calculated *P_m_* and *P_n_* for each of these permuted variants, and then rank ordered the resulting scores. The empirical P-value corresponding to a value of 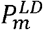 or 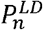 is given by the fraction of permuted variants with higher scores than the given value. We converted these *P*-values into polygenicity/LD-corrected 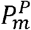 and 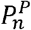 scores by converting each *P*-value into the score it would correspond to under the expected (theoretical) distributions described above.

### Simulation framework

We simulated a realistic matrix of Z-scores *Z* with 100 traits and 800,000 genotyped variants. For non-causal variants, Z-scores for each trait were drawn from an independent Gaussian distribution *N*(0,1). A subset of variants was designated as causal, either pleiotropically or non-pleiotropically. For these causal variants, Z-scores were drawn from a different Gaussian distribution *N*(0, *h*^2^), where *h*^2^ is a parameter representing the per-variant heritability of each trait. Non-pleiotropic variants were selected independently for each trait, while pleiotropic variants were selected globally and each forced to be causal for a specified number of traits *v*. Simulations were run for all combinations of the following parameters: 1) correlation structure: absent or present; 2) proportion of pleiotropic causal variants: 0.1% (800/800,000 variants) or 1% (8,000/800,000 variants); 3) proportion of non-pleiotropic causal variants: 0 (0/800,000 variants), 0.1% (800/800,000 variants), or 1% (8,000/800,000 variants); 4) number of traits involved in horizontal pleiotropy *v*: 10 or 20; 5) per-variant heritability of traits *h*^2^: 0.0002, 0.002, 0. 02, or 0.2. Each scenario was replicated 10,000 times.

### Collection of genome-wide association (GWA) summary statistics datasets

First, we retrieved GWA publicly available summary statistics from 545 continuous traits in 361,194 samples from the UK Biobank (17), and 1,403 binary traits from the same dataset calculated using SAIGE (18,19). We used LD score regression to calculate heritability for each trait, using the liability scale for binary traits, and restricted the sample to only traits with a significant P-value for nonzero heritability after Bonferroni correction. For every pair of traits with correlation coefficient between Z-scores *r*^2^ > 0.8, we additionally removed the member of the pair with lower heritability. This left a total of 372 traits.

Second, we retrieved publicly available genome-wide association (GWA) summary statistics data for 82 complex traits and diseases (31–66) (**Table S9**). For each dataset, we retrieved the appropriate variant annotation (build, rsid, chromosome, position, reference and alternate alleles) and summary statistics (effect size, standard errors, *P*-values and sample size of the study). All variant coordinates (chr, pos) were lifted over to hg19 using the UCSC Genome Browser LiftOver Tool and aligned to the reference and alternate alleles retrieved from the Ensembl variation database. After performing the same pruning of highly correlated phenotypes, we were left with 73 traits and diseases.

Third, we retrieved GWA summary statistics datasets from a GWAS of 453 blood metabolites in 7,824 individuals (67). After performing the same pruning of highly correlated phenotypes, we were left with 430 metabolites.

### Genome-wide pleiotropy study (GWPS)

Using the two components of the pleiotropy score, we can run a genome-wide pleiotropy study (GWPS) which consists of computing two *P*-values for each component of the score (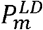 and 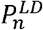) and for all variants genome-wide. We computed the pleiotropy score separately for each of the three datasets described above (372 UK Biobank phenotypes, 73 traits and diseases, and 430 blood metabolites). Additionally, we computed the pleiotropy score on a subset of 372 traits with genome-wide significant heritability as calculated by LD Score Regression (20) (univariate heritability significant after Bonferroni correction). The 372 UK Biobank heritable traits were used for discovery, and the 73 traits and diseases and 430 blood metabolites datasets were used for replication. There was a total of 768,756 variants genotyped across all three datasets.

### Study of polygenicity on horizontal pleiotropy

To study the effect of polygenicity on horizontal pleiotropy, we first estimated the liability-scale heritability of all 372 traits in our UK Biobank dataset using LD score regression, and stratified all traits into four equally-sized classes of heritability, in order to control for the effect of high heritability separate from the effect of high polygenicity. Next, we estimated the polygenicity of the 372 traits using a corrected version of the genomic inflation factor 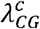 (20). The intercept of LD score regression minus one is an estimator of the mean contribution of confounding bias to the inflation in the test statistics. Therefore, we computed a corrected version of the genomic inflation factor by subtracting the quantity (intercept of LD score regression – 1) from *λ_GC_*. The 372 phenotypes were then ranked according to 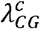 within each heritability class, and grouped into 5 equal-sized bins of about 20 phenotypes each. We then recomputed the LD-corrected pleiotropy score components (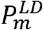 and 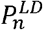) for the subset of phenotypes in each bin. The inflation of the pleiotropy score was measured per bin to represent pleiotropy score inflation as a function of polygenicity.

### Characterization of the pleiotropic variants

We performed various enrichment analyses for the pleiotropy score to characterize the pleiotropic variants using a variety of annotations that could be a direct consequence of horizontal pleiotropy. Each analysis uses the principle of assigning each variant an annotation category and selecting one category as the reference category. Then, for each category, we selected a set of variants from the corresponding reference category with minor allele frequencies matched to those in the query category, and performed a Student’s t-test to test whether the average LD-corrected pleiotropy score (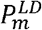 and 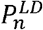) of the variants in each given category is different from the average LD-corrected pleiotropy score of the selected reference variants.

First, we used Ensembl Variant Effect Predictor (21) to classify each variant as noncoding, UTR, nonsynonymous, or coding synonymous, treating noncoding as the reference class. These were complemented by annotations from Roadmap Epigenomics (22). We used the 25-state chromatin state model published by Roadmap Epigenomics to assign each variant 25 scores from 0 to 127, where each score represents the number of epigenomes for which that site is assigned to the corresponding category. We did the same for two specific chromatin marks: the activating mark H3K27ac and the repressive mark H3K27me3. For these annotations, we used a combination of all other categories as a reference set. In other words, the reference set for each category is all variants that are not in that category.

In addition to these molecular annotations, we used expression-related annotations from the Genotype-Tissue Expression project (23). For each variant, we retrieved the number of genes for which the variant is referenced as a *cis* eQTL (expression quantitative trait loci) in any tissue (eGenes), and the number of tissues where the variant is annotated as a *cis* eQTL for any gene (eTissues). We divided variants into bins by number of eGenes (below 10, between 10 and 15, between 15 and 20, and over 20) and eTissues (below 30, between 30 and 35, between 35 and 40, and above 40). The reference set used for these analyses were variants that are not annotated as eQTLs in any gene or tissue.

Finally, we used model organism phenotypes measured by the International Mouse Phenotyping Consortium (IMPC) (68) and the Saccharomyces Cerevisiae Morphological Database (SCMD) (69). To map ortholog genes from IMPC and SCMD to human variants, we used orthology annotations of gene orthologs, and eQTLs from GTEx. Thus, variants annotated as associated with a mouse or yeast phenotype are those that are annotated as *cis* eQTLs in any tissue for a gene whose ortholog in mouse or yeast is associated with that phenotype. The reference set for this analysis was variants annotated as *cis* eQTLs for genes that are not associated with mouse or yeast phenotypes.

### Genome-wide significant pleiotropy loci

To detect loci with a genome-wide significant pleiotropy, we used the LD-corrected two-component pleiotropy score (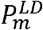 and 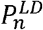) computed on the dataset of 372 heritable traits from UK Biobank described above. We used LD clumping as implemented in PLINK to cluster linked variants, with an *r*^2^ threshold of 0.1, a distance threshold of 100 kb, and *P*-value thresholds of 5 x 10^-8^ for genome-wide significance and 0.05 for nominal significance. The resulting loci were assigned to genes using 1) localization of variants within a gene, as annotated by Ensembl Variant Effect predictor, and 2) annotation as a *cis* eQTL in any tissue, as annotated by GTEx. We submitted the resulting list of genome-wide significant genes to DAVID for enrichment analysis (70–72).

### Permutation-based null model for replication analysis

In general, we should expect only 5% of loci to replicate by chance in each replication dataset; however, it is possible that this number might increase because of polygenicity in the underlying GWAS statistics and the resulting inflation in our pleiotropy score, which may cause substantially more than 5% of the genome to be assigned *P* < 0.05. To correct for this, we performed random permutations of the whitened Z-scores independently for each trait and used these permuted Z-scores to compute our LD-corrected pleiotropy score components (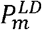 and 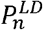). This generates a null expectation that preserves the polygenicity and inflation within each dataset. For both datasets, our null model expected that 5% of loci for 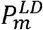 loci and 6% of loci for 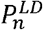 should replicate. The fraction that replicated in the actual data was substantially higher (**Figure 7**).

## Ethics approval and consent to participate

Not applicable.

## Consent for publication

Not applicable.

## Availability of data and material

An R package implementing the HOrizontal Pleiotropy Score (HOPS) method is available on GitHub under the GNU Public License (GPL), at https://github.com/rondolab/HOPS (73). The current version of this repository at the time of submission has been deposited in Zenodo, at https://dx.doi.org/10.5281/zenodo.3462163. The dataset of summary statistics for the 372 medical traits from the UK Biobank and the pleiotropy scores computed from these summary statistics are also available in the same GitHub project at https://github.com/rondolab/HOPS/tree/master/data. The summary statistics for 430 blood metabolites are available from the original publication where this dataset was reported (61), and the summary statistics for 73 human traits and diseases are available from the original publications where they were reported, as cited in **Additional File 1: Table S3**.

## Authors’ contributions

D.M.J. and M.V. contributed to study conception, data analysis, interpretation of the results and drafting of the manuscript. R.D. contributed to study conception, interpretation of the results and critical revision of the manuscript.

## Acknowledgments

We thank the various genome-wide association consortia as well as Dr. Benjamin Neale’s and Dr. Seunggeun Lee’s group for generously sharing genome-wide association summary statistics for phenome-wide association scan in UK biobank.

## Additional Files

**Additional File 1. Supplementary tables and figures.**

Word document (.docx) containing supplementary figures S1-S8, and supplementary tables S1-S4.

**Additional File 2. Functional enrichment analysis of pleiotropy score after applying polygenicity correction.**

Excel spreadsheet (.xlsx) showing the equivalent of **Table 1** using the LD/polygenicity-corrected scores (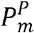 and 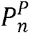) instead of the LD-corrected scores (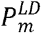 and 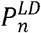)

**Additional File 3. DAVID enrichment analysis of high-pleiotropy genes.**

Excel spreadsheet (.xlsx) showing the results of the DAVID enrichment analysis described in the text.

