## Additional File 1 for "HOPS: a quantitative score reveals pervasive horizontal pleiotropy in human genetic variation is driven by extreme polygenicity of human traits and diseases"

### Power to detect pleiotropy in simulations

**Fig. S1: Effect of LD quantile on two-component pleiotropy score.** Dots show simulated $P_{m}$ (left) and $P_{n}$ (right), both with and without LD-correction, stratified by the simulated level of LD. In both cases, the highest quantile of LD shows substantially inflated pleiotropy score without LD-correction, but applying the LD correction reduces the pleiotropy score to the level seen when there is no LD.

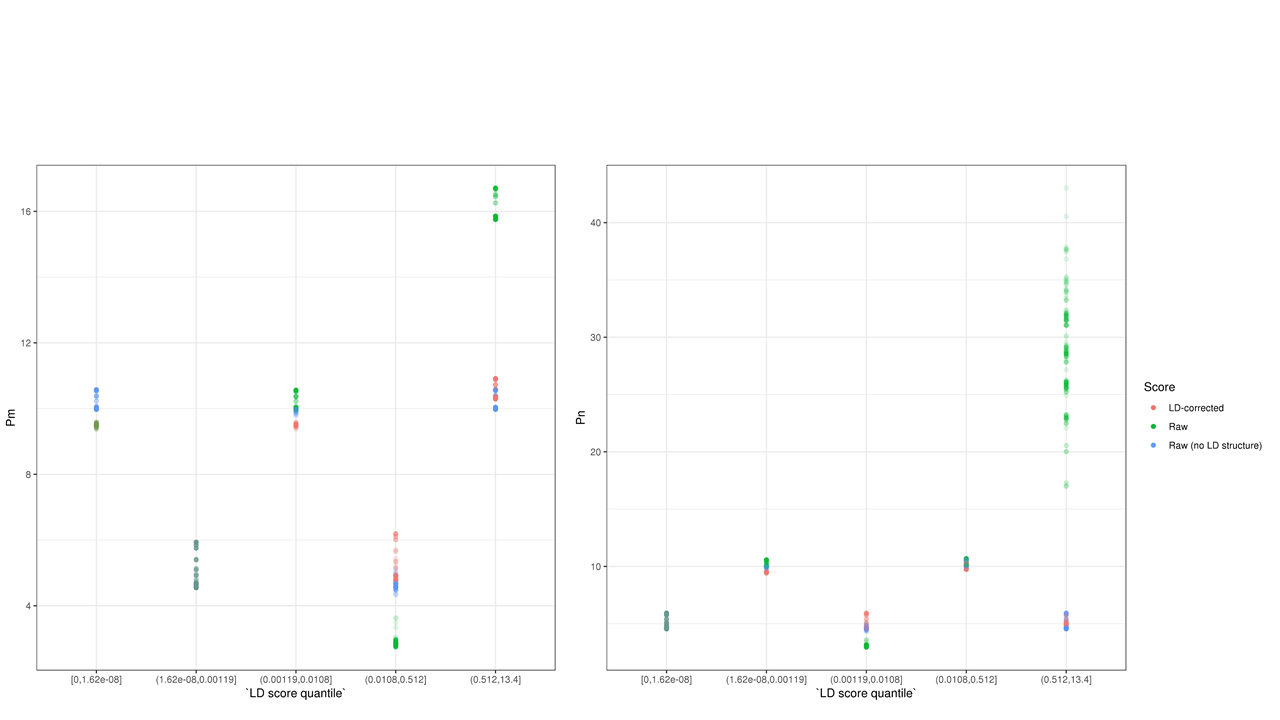

**Fig. S2: Simulation study showing power of two-component pleiotropy score with increased number of traits effected by pleiotropy.**

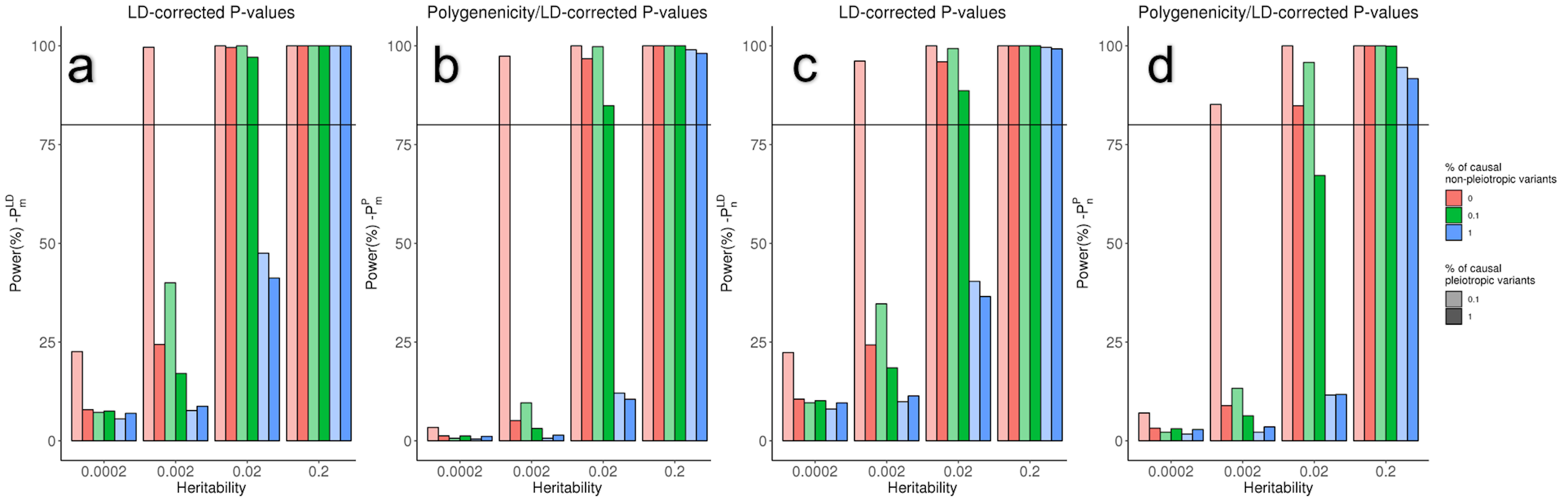

Black line shows 80% power. Simulations were run for both $P_{m}^{LD}$ (left) and $P_{n}^{LD}$ (right), and both without correction for polygenicity (a,c) and with the correction (b,d), with per-variant heritability ranging from 0.0002 to 0.2, proportion of non-pleiotropic causal loci ranging from 0 to 1%, and proportion of pleiotropic causal loci ranging from 0.1% to 1%. This figure shows a scenario where each pleiotropic variant affects 20 traits, as opposed to the main text figure (**Figure 2**) which shows each pleiotropic variant affecting 10 traits.

**Fig. S3: Simulation study showing power of two-component pleiotropy score with correlation structure.**

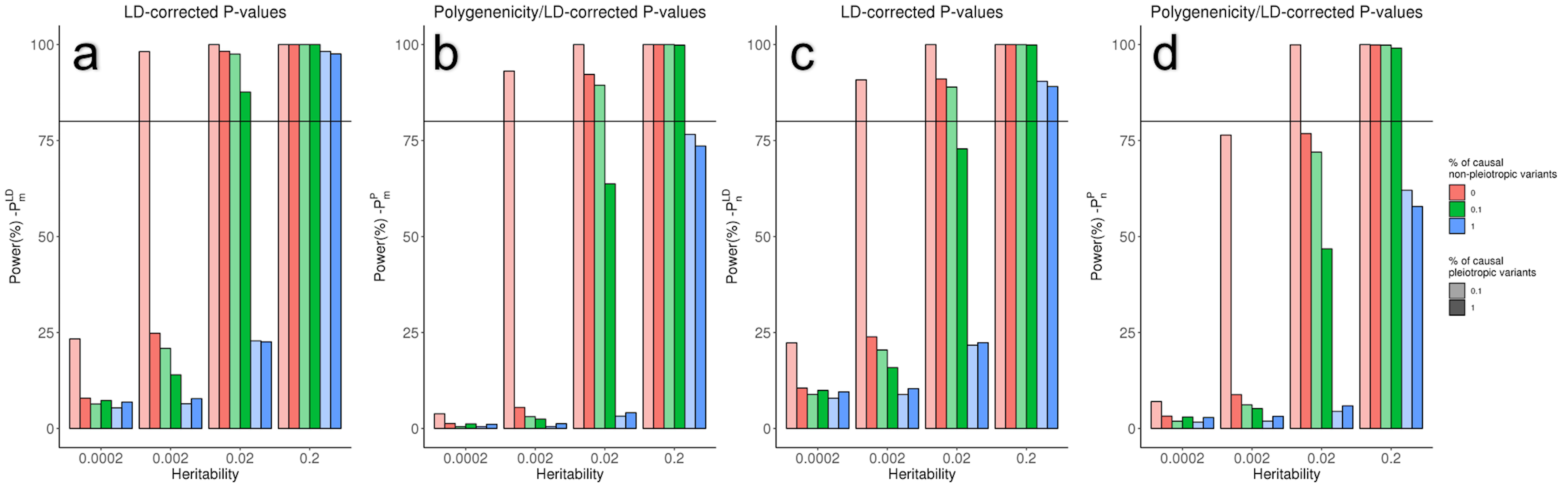

Black line shows 80% power. Simulations were run for both $P_{m}^{LD}$ (left) and $P_{m}^{LD}$ (right), and both without correction for polygenicity (a,c) and with the correction (b,d), with per-variant heritability ranging from 0.0002 to 0.2, proportion of non-pleiotropic causal loci ranging from 0 to 1%, and proportion of pleiotropic causal loci ranging from 0.1% to 1%. This figure shows a scenario where traits have correlations between them, as opposed to the main text figure (**Figure 2**) which shows uncorrelated traits.

### Genome-Wide Pleiotropy Study (GWPS) reveals pervasive pleiotropy

Fig. S4: Quantile-quantile plots (Q-Q plot) of the genome-wide pleiotropy study (GWPS) for the 372 medical traits from the UK Biobank.

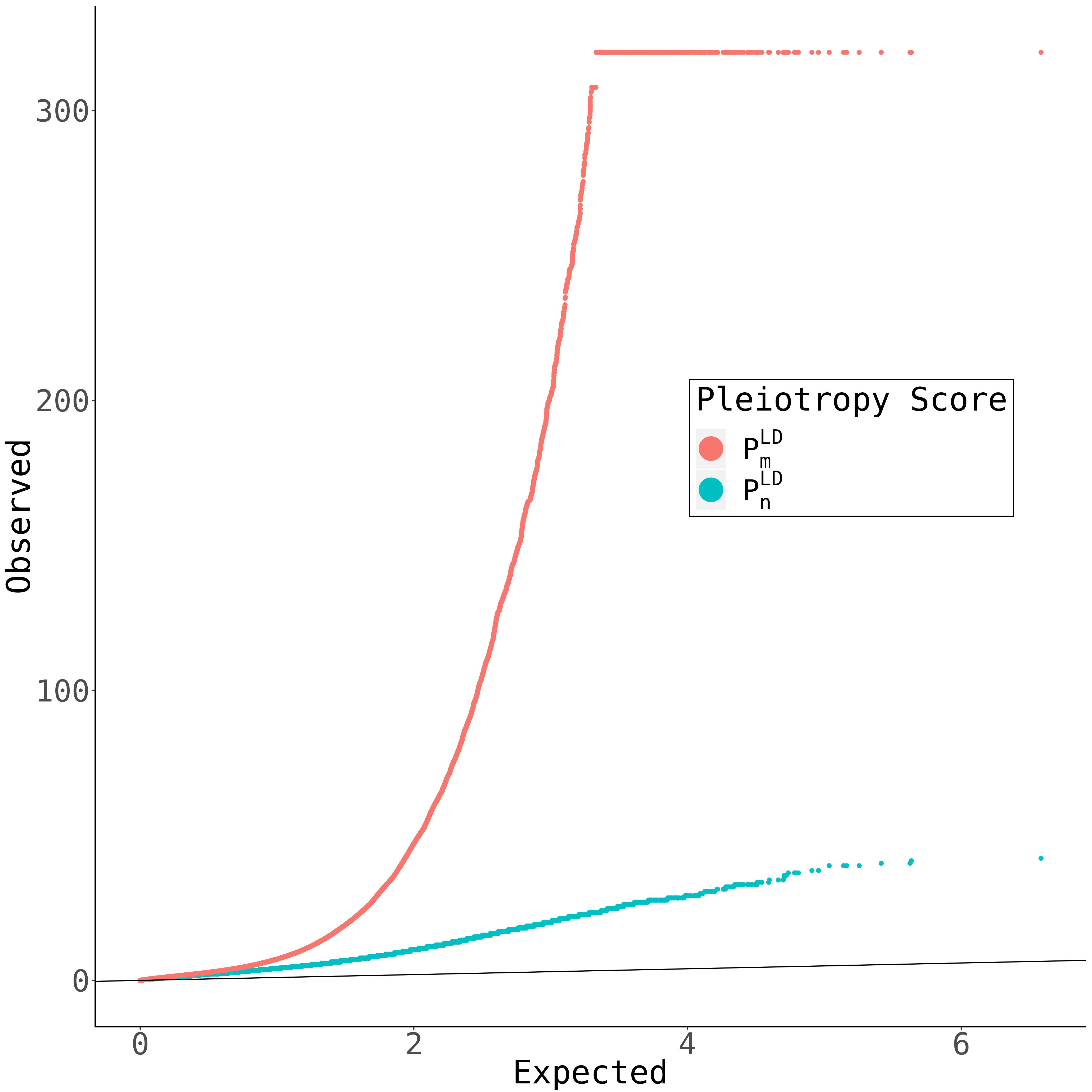

In each plot the x-axis represent the expected distribution of *P-values*, and the y-axis the observed *P-values* for $P_{m}^{LD}$ (red dots) and $P_{n}^{LD}$ (blue dots). The line shows the observed *P-values* under the null hypothesis. -log_10_ transformed *P-values* are represented.

**Fig. S5: Manhattan plot of the genome-wide pleiotropy study (GWPS) using the two-component pleiotropy score on the 372 heritable medical traits from the UK Biobank.**

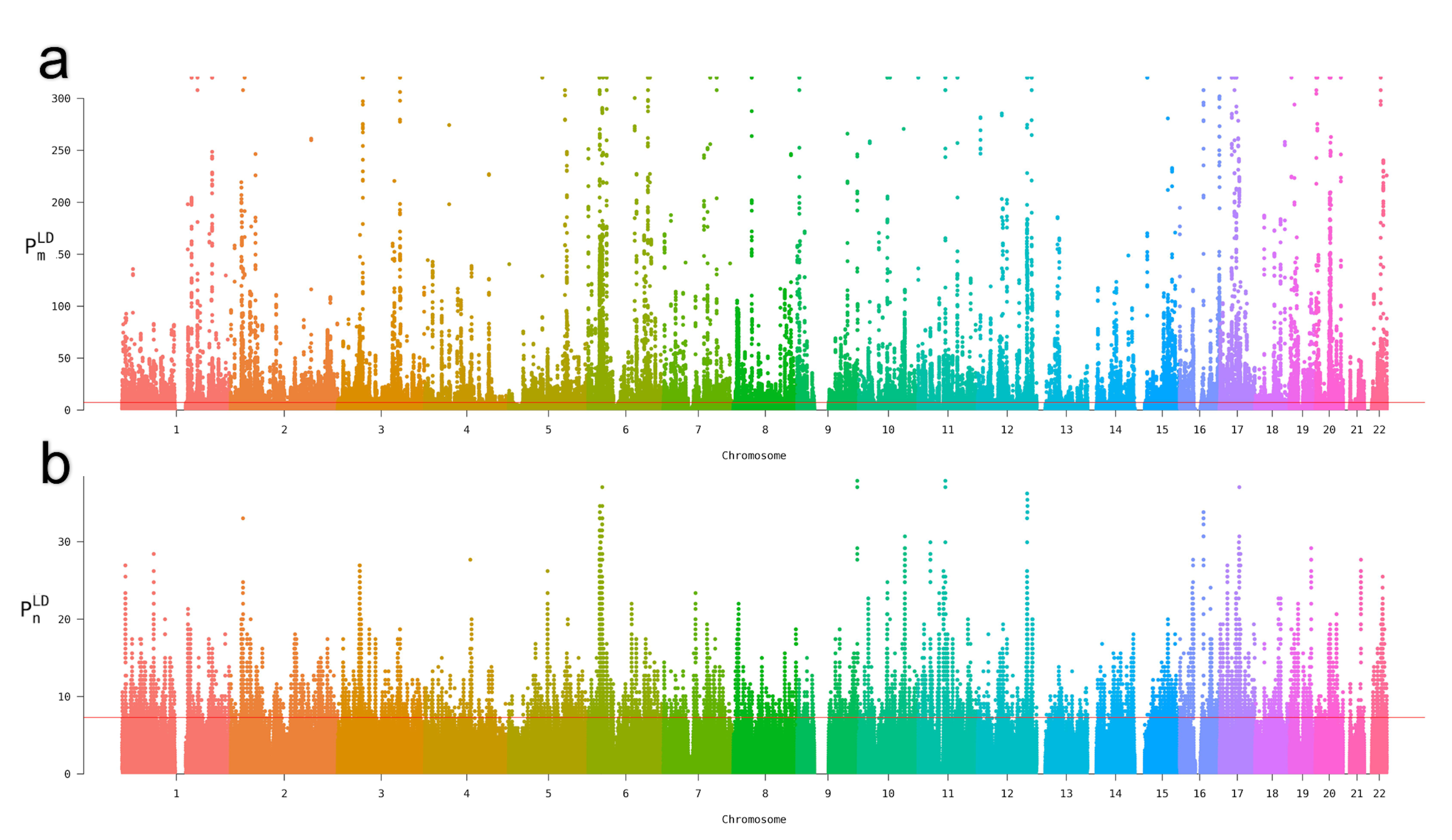

The results of the GWPS are provided for the two components of the pleiotropy score, $P_{m}^{LD}$ (top) and $P_{n}^{LD}$ (bottom) on the 372 heritable medical traits from the UK Biobank. Each dot representing a variant is colored by chromosome and ordered by position on the x axis. -log_10_ transformed *P-values* are represented on the y axis. The red line represents the genome-wide significant cut-off of 5 x 10^-8^

**Fig. S6: Scatterplot of the two-component pleiotropy score (left:** $\boldsymbol{P}_{\boldsymbol{m}}^{\boldsymbol{LD}}$**, right:** $\boldsymbol{P}_{\boldsymbol{n}}^{\boldsymbol{LD}}$**) between two decorrelation procedures, using all variants (All) or an independent subset (Pruned) to estimate the phenotype-correlation matrix.**

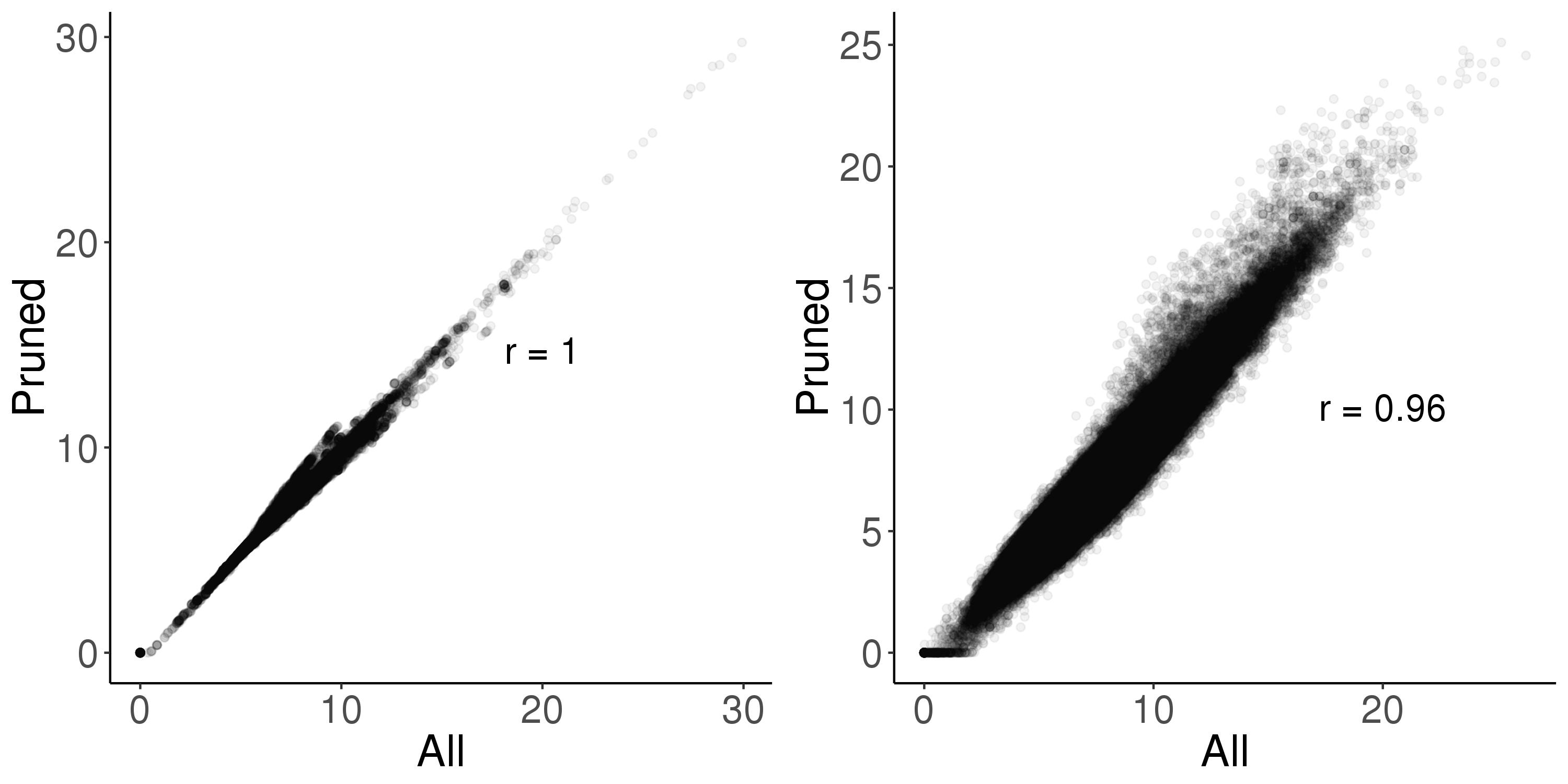

### Pleiotropy is driven by polygenicity

Fig. S7: Quantile-quantile plots (Q-Q plot) of the genome-wide pleiotropy study (GWPS) for the 372 medical traits from the UK Biobank after applying the correction for polygenicity.

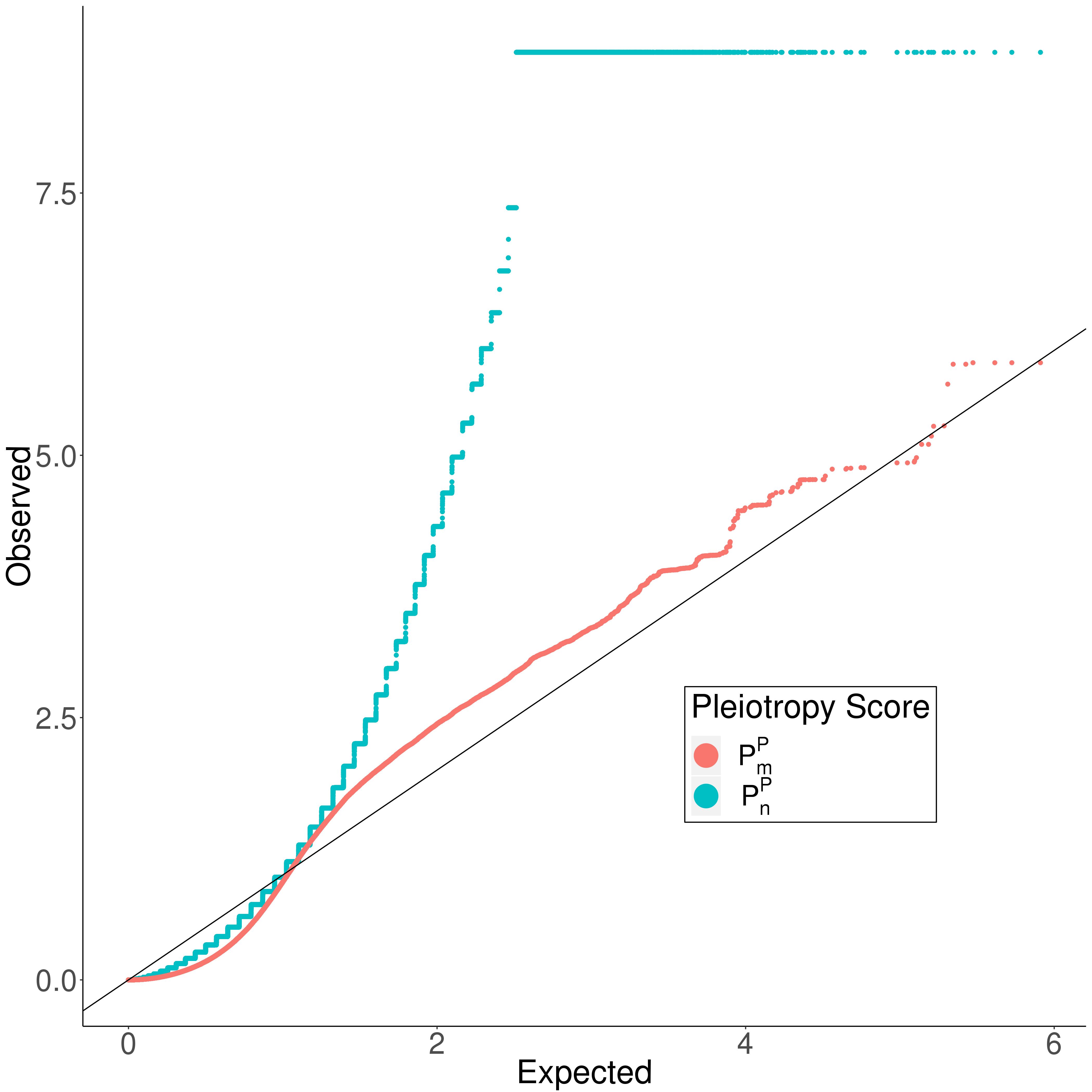

In each plot the x-axis represent the expected distribution of *P-values*, and the y-axis the observed *P-values* for $P_{m}^{P}$ (red dots) and $P_{n}^{P}$ (blue dots). The line shows the observed *P-values* under the null hypothesis. -log_10_ transformed *P-values* are represented.

**Fig. S8: Manhattan plot of the genome-wide pleiotropy study (GWPS) using the two-component polygenicity/LD-corrected pleiotropy score on the 372 heritable medical traits from the UK Biobank after applying the correction for polygenicity.**

The results of the GWPS are provided for the two components of the polygenicity/LD-corrected pleiotropy score, $P_{m}^{P}$ (top) and $P_{n}^{P}$ (bottom) on the 372 heritable medical traits from the UK Biobank. Each dot representing a variant is colored by chromosome and ordered by position on the x axis. -log_10_ transformed *P-values* are represented on the y axis. The red line in panel b represents the genome-wide significant cut-off of 5 x 10^-8^; no variants rise to the level of genome-wide significance in $P_{m}^{P}$ (panel a).

Table S1: Effect of polygenicity on pleiotropy score

|  |  |  |  | **% genome-wide significant variants** | | | **Inflation factor** $\lambda_{GC}$ **of the pleiotropy score** | | | **Mann-Whitney test (greater)** | | | **Mann-Whitney test (less)** | |
| --- | --- | --- | --- | --- | --- | --- | --- | --- | --- | --- | --- | --- | --- | --- |
| **Range of** $\boldsymbol{h}^{\boldsymbol{2}}$ | **Range of** $\lambda_{GC}^{c}$ | **Average** $\boldsymbol{h}^{\boldsymbol{2}}$ | **# traits** | $\boldsymbol{P}_{\boldsymbol{m}}^{\boldsymbol{LD}}$ | $\boldsymbol{P}_{\boldsymbol{n}}^{\boldsymbol{LD}}$ | $\boldsymbol{P}_{\boldsymbol{m}}^{\boldsymbol{LD}}$ | | $\boldsymbol{P}_{\boldsymbol{n}}^{\boldsymbol{LD}}$ | $\boldsymbol{P}_{\boldsymbol{m}}^{\boldsymbol{LD}}$ | | $\boldsymbol{P}_{\boldsymbol{n}}^{\boldsymbol{LD}}$ | $\boldsymbol{P}_{\boldsymbol{m}}^{\boldsymbol{LD}}$ | | $\boldsymbol{P}_{\boldsymbol{n}}^{\boldsymbol{LD}}$ |
| 0.76%-4.3% | 1.029-1.046 | 2.942 | 28 | 0.0010 | 0 | 0.9685 | | 0.6623 | 1 | | 1 | 8.63x10^-19^ | | 0 |
| 0.76%-4.3% | 1.046-1.063 | 2.691 | 28 | 0.0012 | 0 | 1.1153 | | 0.6623 | 1.08x10^-291^ | | 1 | 1 | | 0 |
| 0.76%-4.3% | 1.063-1.091 | 2.620 | 28 | 0.0116 | 0.0003 | 1.3596 | | 0.6623 | 0 | | 1 | 1 | | 0 |
| 0.76%-4.3% | 1.091-1.146 | 2.608 | 28 | 0.0005 | 0 | 1.2836 | | 0.6623 | 0 | | 1 | 1 | | 0 |
| 0.76%-4.3% | 1.146-1.251 | 3.548 | 27 | 0.0594 | 0.0005 | 1.6868 | | 0.6623 | 0 | | 1 | 1 | | 3.11x10^-40^ |
| 4.3%-7.27% | 1.033-1.053 | 5.364 | 28 | 0.0545 | 0.0001 | 1.2872 | | 0.6623 | 0 | | 1 | 1 | | 0 |
| 4.3%-7.27% | 1.053-1.079 | 6.048 | 28 | 0.2564 | 0.0063 | 2.1483 | | 0.6623 | 0 | | 0 | 1 | | 1 |
| 4.3%-7.27% | 1.079-1.118 | 5.650 | 28 | 1.1707 | 0.1471 | 5.4103 | | 3.4573 | 0 | | 0 | 1 | | 1 |
| 4.3%-7.27% | 1.118-1.273 | 5.505 | 28 | 5.4601 | 0.8052 | 5.9886 | | 3.4573 | 0 | | 0 | 1 | | 1 |
| 4.3%-7.27% | 1.273-1.38 | 6.117 | 27 | 5.9312 | 1.2484 | 4.8754 | | 3.4573 | 0 | | 0 | 1 | | 1 |
| 7.27%-12.79% | 1.031-1.056 | 9.830 | 28 | 0.0055 | 0 | 1.1411 | | 0.6623 | 0 | | 1 | 1 | | 0 |
| 7.27%-12.79% | 1.056-1.093 | 9.448 | 28 | 0.0102 | 0 | 1.2913 | | 0.6623 | 0 | | 1 | 1 | | 0 |
| 7.27%-12.79% | 1.093-1.157 | 9.921 | 28 | 0.0065 | 0 | 1.4285 | | 0.6623 | 0 | | 1 | 1 | | 0 |
| 7.27%-12.79% | 1.157-1.268 | 9.981 | 28 | 0.0236 | 0.0009 | 1.4573 | | 0.6623 | 0 | | 1 | 1 | | 0 |
| 7.27%-12.79% | 1.268-1.587 | 9.824 | 27 | 0.0080 | 0.0008 | 1.6934 | | 0.6623 | 0 | | 0.9996 | 1 | | 0.0004 |
| 12.79%-42.19% | 1.028-1.138 | 16.434 | 28 | 0.0231 | 0 | 1.2148 | | 0.6623 | 0 | | 1 | 1 | | 0 |
| 12.79%-42.19% | 1.138-1.254 | 22.421 | 28 | 0.0405 | 0.0001 | 1.2222 | | 0.6623 | 0 | | 1 | 1 | | 0 |
| 12.79%-42.19% | 1.254-1.415 | 21.446 | 28 | 0.0628 | 0.0020 | 1.4249 | | 0.6623 | 0 | | 1 | 1 | | 0 |
| 12.79%-42.19% | 1.415-1.912 | 20.373 | 28 | 0.1056 | 0.0035 | 1.5504 | | 0.6623 | 0 | | 1 | 1 | | 3.51x10^-36^ |
| 12.79%-42.19% | 1.912-2.045 | 23.097 | 27 | 0.1506 | 0.0103 | 1.9366 | | 0.6623 | 0 | | 0 | 1 | | 1 |

### Genome-wide pleiotropy study identifies novel biological loci

Table S2: Counts of genome-wide significant variants for the pleiotropy score.

|  |  | $P_{n}^{LD}$ | | |
| --- | --- | --- | --- | --- |
|  |  | *P* > 0.05 | 5 x 10^-8^ < *P* < 0.05 | P < 5 x 10^-8^ |
| $P_{m}^{LD}$ | *P* > 0.05 | 272,396 | 33,998 | 0 |
|  | 5 x 10^-8^ < *P* < 0.05 | 113,505 | 265,662 | 354 |
|  | P < 5 x 10^-8^ | 6,529 | 56,027 | 18,587 |

### Pleiotropic loci replicate in independent GWAS datasets

Table S3: Summary of publicly available genome-wide association summary statistic datasets used in the present study.

| Category | Name | Consortium | Link | Reference |
| --- | --- | --- | --- | --- |
| CAD | Total cholesterol | Global Lipids Genetics | [Link](http://csg.sph.umich.edu/abecasis/public/lipids2013/jointGwasMc_TC.txt.gz) | (1) |
| CAD | Low-density lipoprotein cholesterol | Global Lipids Genetics | [Link](http://csg.sph.umich.edu/abecasis/public/lipids2013/jointGwasMc_LDL.txt.gz) | (1) |
| CAD | High-density lipoprotein cholesterol | Global Lipids Genetics | [Link](http://csg.sph.umich.edu/abecasis/public/lipids2013/jointGwasMc_HDL.txt.gz) | (1) |
| CAD | Triglycerides | Global Lipids Genetics | [Link](http://csg.sph.umich.edu/abecasis/public/lipids2013/jointGwasMc_TG.txt.gz) | (1) |
| CAD | Coronary artery disease | CARDIoGRAMplusC4D | [Link](ftp://ftp.sanger.ac.uk/pub/cardiogramplusc4d/2015/cad.additive.Oct2015.pub.zip) | (2) |
| Adiposity | Body mass index | GIANT | [Link](https://www.broadinstitute.org/collaboration/giant/images/1/15/SNP_gwas_mc_merge_nogc.tbl.uniq.gz) | (3) |
| Adiposity | Waist circumference | GIANT | [Link](https://www.broadinstitute.org/collaboration/giant/images/5/57/GIANT_2015_WC_COMBINED_EUR.txt.gz) | (4) |
| Adiposity | Waist circumference (Adj BMI) | GIANT | [Link](https://www.broadinstitute.org/collaboration/giant/images/7/73/GIANT_2015_WCadjBMI_COMBINED_EUR.txt.gz) | (4) |
| Adiposity | Hip circumference | GIANT | [Link](https://www.broadinstitute.org/collaboration/giant/images/e/e4/GIANT_2015_HIP_COMBINED_EUR.txt.gz) | (4) |
| Adiposity | Hip circumference (Adj BMI) | GIANT | [Link](https://www.broadinstitute.org/collaboration/giant/images/5/52/GIANT_2015_HIPadjBMI_COMBINED_EUR.txt.gz) | (4) |
| Adiposity | Waist hip ratio | GIANT | [Link](https://www.broadinstitute.org/collaboration/giant/images/5/54/GIANT_2015_WHR_COMBINED_EUR.txt.gz) | (4) |
| Adiposity | Waist hip ratio (Adj BMI) | GIANT | [Link](https://www.broadinstitute.org/collaboration/giant/images/e/eb/GIANT_2015_WHRadjBMI_COMBINED_EUR.txt.gz) | (4) |
| Adiposity | Height | GIANT | [Link](https://www.broadinstitute.org/collaboration/giant/images/0/01/GIANT_HEIGHT_Wood_et_al_2014_publicrelease_HapMapCeuFreq.txt.gz) | (5) |
| Adiposity | Adiponectin | ADIPOGen | [Link](https://www.mcgill.ca/genepi/files/genepi/adipogen.discovery.eur_.meta_.public.release.part1_.txt) | (6) |
| T2D | Type-2 diabetes | DIAGRAM | [Link](http://diagram-consortium.org/downloads.html) | (7) |
| T2D | Systolic blood pressure | ICBP | Request | (8) |
| T2D | Diastolic blood pressure | ICBP | Request | (8) |
| T2D | Fasting insulin | MAGIC | [Link](ftp://ftp.sanger.ac.uk/pub/magic/MAGIC_ln_FastingInsulin.txt) | (9) |
| T2D | HOMA IR | MAGIC | [Link](ftp://ftp.sanger.ac.uk/pub/magic/MAGIC_ln_HOMA-IR.txt) | (10) |
| T2D | Fasting insulin interaction | MAGIC | [Link](ftp://ftp.sanger.ac.uk/pub/magic/MAGIC_Manning_et_al_lnFastingInsulin_Interaction.txt.gz) | (11) |
| T2D | Fasting proinsulin | MAGIC | [Link](ftp://ftp.sanger.ac.uk/pub/magic/MAGIC_ln_fastingProinsulin.txt.gz) | (9) |
| T2D | Area under the curve - insulin/glucose | MAGIC | [Link](ftp://ftp.sanger.ac.uk/pub/magic/MAGIC_InsulinSecretion_data_release_May2014.zip) | (12) |
| T2D | Area under the curve - insulin | MAGIC | [Link](ftp://ftp.sanger.ac.uk/pub/magic/MAGIC_InsulinSecretion_data_release_May2014.zip) | (12) |
| T2D | Corrected insulin response | MAGIC | [Link](ftp://ftp.sanger.ac.uk/pub/magic/MAGIC_InsulinSecretion_data_release_May2014.zip) | (12) |
| T2D | Corrected insulin response / insulin sensitivity | MAGIC | [Link](ftp://ftp.sanger.ac.uk/pub/magic/MAGIC_InsulinSecretion_data_release_May2014.zip) | (12) |
| T2D | DI | MAGIC | [Link](ftp://ftp.sanger.ac.uk/pub/magic/MAGIC_InsulinSecretion_data_release_May2014.zip) | (12) |
| T2D | Incretin 30 min | MAGIC | [Link](ftp://ftp.sanger.ac.uk/pub/magic/MAGIC_InsulinSecretion_data_release_May2014.zip) | (12) |
| T2D | Insulin 30 min (Adj BMi) | MAGIC | [Link](ftp://ftp.sanger.ac.uk/pub/magic/MAGIC_InsulinSecretion_data_release_May2014.zip) | (12) |
| T2D | Insulin 30 min | MAGIC | [Link](ftp://ftp.sanger.ac.uk/pub/magic/MAGIC_InsulinSecretion_data_release_May2014.zip) | (12) |
| T2D | Insulin sensitivity | MAGIC | [Link](ftp://ftp.sanger.ac.uk/pub/magic/MAGIC_InsulinSecretion_data_release_May2014.zip) | (12) |
| T2D | Fasting glucose | MAGIC | [Link](ftp://ftp.sanger.ac.uk/pub/magic/MAGIC_ln_FastingGlucose.txt) | (9) |
| T2D | HOMA B | MAGIC | [Link](ftp://ftp.sanger.ac.uk/pub/magic/MAGIC_ln_HOMA-B.txt) | (10) |
| T2D | Fasting glucose interaction | MAGIC | [Link](ftp://ftp.sanger.ac.uk/pub/magic/MAGIC_Manning_et_al_FastingGlucose_Interaction.txt.gz) | (11) |
| T2D | Two-hour glucose | MAGIC | [Link](ftp://ftp.sanger.ac.uk/pub/magic/MAGIC_2hrGlucose_AdjustedForBMI.txt) | (9) |
| T2D | Glycated Hemoglobin A1C | MAGIC | [Link](ftp://ftp.sanger.ac.uk/pub/magic/MAGIC_HbA1C.txt.gz) | (9) |
| CKD | Chronic kidney disease | CKDGEN | [Link](http://ckdgen.imbi.uni-freiburg.de/files/Pattaro2016/formatted_round3meta_CKD_overall_IV_2GC_b36_MAFget005_Nget50_20120725_b37.csv.gz) | (13) |
| CKD | Estimated glomerular filtration rate (creatinine) | CKDGEN | [Link](http://ckdgen.imbi.uni-freiburg.de/files/Pattaro2016/formatted_round3meta_eGFRcrea_overall_IV_2GC_b36_MAFget005_Nget50_20120705_b37.csv.gz) | (13) |
| CKD | Estimated glomerular filtration rate (cystatin) | CKDGEN | [Link](http://ckdgen.imbi.uni-freiburg.de/files/Pattaro2016/formatted_round3meta_eGFRcys_overall_IV_2GC_b36_MAFget005_Nget50_20120725_b37.csv.gz) | (13) |
| CKD | Urate | GUGC | Request | (14) |
| Inflammatory Markers | C-reactive protein | CHARGE | Request | (15) |
| Inflammatory Markers | Lipoprotein a | Framingham Heart Study | Request |  |
| Inflammatory Diseases | Rheumatoid arthritis | - | [Link](http://plaza.umin.ac.jp/~yokada/datasource/files/GWASMetaResults/RA_GWASmeta_European_v2.txt.gz) | (16) |
| Inflammatory Diseases | Crohn's disease | IIBDGC | [Link](ftp://ftp.sanger.ac.uk/pub/consortia/ibdgenetics/cd-meta.txt.gz) | (17) |
| Inflammatory Diseases | Inflammatory bowel disease | IIBDGC | [Link](ftp://ftp.sanger.ac.uk/pub/consortia/ibdgenetics/iibdgc-trans-ancestry-summary-stats.tar) | (17) |
| Inflammatory Diseases | Ulcerative colitis | IIBDGC | [Link](ftp://ftp.sanger.ac.uk/pub/consortia/ibdgenetics/ucmeta-sumstats.txt.gz) | (17) |
| Inflammatory Diseases | Gout | GUGC | Request | (14) |
| Inflammatory Diseases | Alzheimer's disease | IGAP | [Link](http://www.pasteur-lille.fr/en/recherche/u744/igap/igap_download.php) | (18) |
| Psychiatric Disorders | Bipolar disorder | PGC | [Link](https://www.med.unc.edu/pgc/results-and-downloads/data-use-agreement-forms/BIP_data_download_agreement) | (19) |
| Psychiatric Disorders | Schizophrenia | PGC | [Link](https://www.med.unc.edu/pgc/results-and-downloads/data-use-agreement-forms/SCZSNP_data_download_agreement) | (20) |
| Psychiatric Disorders | Attention deficit hyperactivity disorder | PGC | [Link](https://www.med.unc.edu/pgc/results-and-downloads/data-use-agreement-forms/CrossADHD_data_download_agreement) | (21) |
| Psychiatric Disorders | Autism spectrum disorder | PGC | [Link](https://www.med.unc.edu/pgc/results-and-downloads/data-use-agreement-forms/CrossASD_data_download_agreement) | (22) |
| Psychiatric Disorders | Eating disorder | PGC | [Link](https://www.med.unc.edu/pgc/results-and-downloads/data-use-agreement-forms/ED_data_download_agreement) | (22) |
| Psychiatric Disorders | Major depressive disorder | PGC | [Link](https://www.med.unc.edu/pgc/results-and-downloads/data-use-agreement-forms/MDD_data_download_agreement) | (23) |
| Brain Volume | Intracranial volume | ENIGMA | [Link](http://enigma.ini.usc.edu/research/download-enigma-gwas-results/data-agreement-for-enigma2-download/) | (24) |
| Brain Volume | Mean accumbens | ENIGMA | [Link](http://enigma.ini.usc.edu/research/download-enigma-gwas-results/data-agreement-for-enigma2-download/) | (24) |
| Brain Volume | Mean amygdala | ENIGMA | [Link](http://enigma.ini.usc.edu/research/download-enigma-gwas-results/data-agreement-for-enigma2-download/) | (24) |
| Brain Volume | Mean caudate | ENIGMA | [Link](http://enigma.ini.usc.edu/research/download-enigma-gwas-results/data-agreement-for-enigma2-download/) | (24) |
| Brain Volume | Mean hippocampus | ENIGMA | [Link](http://enigma.ini.usc.edu/research/download-enigma-gwas-results/data-agreement-for-enigma2-download/) | (24) |
| Brain Volume | Mean pallidum | ENIGMA | [Link](http://enigma.ini.usc.edu/research/download-enigma-gwas-results/data-agreement-for-enigma2-download/) | (24) |
| Brain Volume | Mean putamen | ENIGMA | [Link](http://enigma.ini.usc.edu/research/download-enigma-gwas-results/data-agreement-for-enigma2-download/) | (24) |
| Brain Volume | Mean thalamus | ENIGMA | [Link](http://enigma.ini.usc.edu/research/download-enigma-gwas-results/data-agreement-for-enigma2-download/) | (24) |
| Red blood cell | Hemoglobin | HaemGen | [Link](https://ega-archive.org/studies/EGAS00000000132) | (25) |
| Red blood cell | Mean cell hemoglobin | HaemGen | [Link](https://ega-archive.org/studies/EGAS00000000132) | (25) |
| Red blood cell | Mean corpuscular hemoglobin concentration | HaemGen | [Link](https://ega-archive.org/studies/EGAS00000000132) | (25) |
| Red blood cell | Mean cell volume | HaemGen | [Link](https://ega-archive.org/studies/EGAS00000000132) | (25) |
| Red blood cell | Packed cell volume | HaemGen | [Link](https://ega-archive.org/studies/EGAS00000000132) | (25) |
| Red blood cell | Red blood cell count | HaemGen | [Link](https://ega-archive.org/studies/EGAS00000000132) | (25) |
| Reproductive Genetics | Age Menarche | ReproGen | [Link](http://www.reprogen.org/Menarche_Nature2014_GWASMetaResults_17122014.zip) | (26) |
| Reproductive Genetics | Age Menopause | ReproGen | [Link](http://www.reprogen.org/Menopause_HapMap2_DayNG2015_18112015.txt.gz) | (27) |
| Early Birth | Birth length | EGG | [Link](http://egg-consortium.org/downloads/EGG-GWAS-BL.txt.gz) | (28) |
| Early Birth | Birth weight | EGG | [Link](http://egg-consortium.org/downloads/EGG_BW2_DISCOVERY.txt.gz) | (29) |
| Early Birth | Childhood BMI | EGG | [Link](http://egg-consortium.org/downloads/EGG_BMI_HapMap_DISCOVERY.txt.gz) | (30) |
| Early Birth | Childhood obesity | EGG | [Link](http://egg-consortium.org/downloads/EGG_Obesity_Meta_Analysis_1.txt.gz) | (31) |
| Early Birth | Head circumference | EGG | [Link](http://egg-consortium.org/downloads/EGG_HC_DISCOVERY.txt.gz) | (32) |
| Early Birth | Pubertal growth | EGG | [Link](http://egg-consortium.org/downloads/Pubertal_growth_10F_12M_combined.txt.gz) | (33) |
| Early Birth | Total growth across pubertal growth period | EGG | [Link](http://egg-consortium.org/downloads/Pubertal_growth_PGF_PGM_combined.txt.gz) | (33) |
| Early Birth | Total growth in late adolescent | EGG | [Link](http://egg-consortium.org/downloads/Pubertal_growth_PTF_PTM_combined.txt.gz) | (33) |
| Social Science | Number of years of education | SSGAC | [Link](http://ssgac.org/documents/SSGAC_Rietveld2013.zip) | (34) |
| Smoking History | Number of cigarettes per day | TGC | [Link](https://www.med.unc.edu/pgc/files/resultfiles/tag.cpd.tbl.gz) | (35) |
| Smoking History | Age of onset smoking | TGC | [Link](https://www.med.unc.edu/pgc/files/resultfiles/tag.logonset.tbl.gz) | (35) |
| Smoking History | Ever smoked | TGC | [Link](https://www.med.unc.edu/pgc/files/resultfiles/tag.evrsmk.tbl.gz) | (35) |
| Smoking History | Former smoker | TGC | [Link](https://www.med.unc.edu/pgc/files/resultfiles/tag.former.tbl.gz) | (35) |
| Metabolites | 453 metabolites | Shin et al. | [Link](http://mips.helmholtz-muenchen.de/proj/GWAS/gwas/index.php?task=download) | (36) |
| UK biobank | 2,419 phenotypes | Neale lab | [Link](http://www.nealelab.is/blog/2017/7/19/rapid-gwas-of-thousands-of-phenotypes-for-337000-samples-in-the-uk-biobank) |  |

### Pleiotropy is correlated with specific complex traits and diseases

Table S4: Sensitivity analysis of the pleiotropy score according to the maximum sample size of summary statistics datasets.

|  | **Maximum sample size** | $\lambda_{GC}$ | **Correlation with** $P_{m}^{LD}$ | **Correlation with** $P_{n}^{LD}$ |
| --- | --- | --- | --- | --- |
| BMI (All GIANT) | 339,224 | 1.171 | 0.192 | 0.201 |
| BMI (Men) | 152,893 | 1.072 | 0.138 | 0.141 |
| BMI (Women) | 171,976 | 1.073 | 0.147 | 0.157 |
| CAD (CARDIoGRAM) | 84,264 | 1.105 | 0.071 | 0.070 |
| CAD (CARDIoGRAM+C4D) | 184,305 | 1.062 | 0.124 | 0.119 |

BMI: Body mass index; CAD: Coronary artery disease.
